# Ontogenetic Patterns in Juvenile Blue Crab density: Effects of Habitat and Turbidity in a Chesapeake Bay Tributary

**DOI:** 10.1101/2023.06.05.543724

**Authors:** A. Challen Hyman, Grace S. Chiu, Michael S. Seebo, Alison Smith, Gabrielle G. Saluta, Romuald N. Lipcius

## Abstract

Nursery habitats are characterized by favorable conditions for juveniles, such as higher food availability and lower predation risk, and disproportionately contribute more individuals per unit area to adult segments of the population compared to other habitats. However, nursery habitat inference is complicated by changes in habitat preferences with ontogeny; individuals in early-life stages frequently inhabit different habitats than older juveniles or adults. In this mensurative field experiment, we modeled the density of two juvenile blue crab, *Callinectes sapidus*, size classes based on carapace width (CW) across multiple habitats at various locations within an estuarine seascape during the blue crab recruitment season. We examined four habitat types—unstructured sand, seagrass meadows, salt marsh edge (SME), and shallow detrital habitat (SDH). Results indicated that densities of small juvenile blue crabs (*≤*15 mm CW) were highest in seagrass, whereas densities of larger juveniles (16–30 mm CW) were highest in SME. Densities of large juveniles in SME were also greater than those of small juveniles, suggesting possible secondary dispersal to SME by small juveniles after settlement and recruitment in seagrass. Turbidity was positively correlated with densities of both size classes, although our model did not address whether this was due to top-down (refuge) or bottom-up (food availability) mechanisms. Observed patterns in size-specific habitat utilization may result from changing requirements of juvenile blue crabs with size, as animals minimize mortality-to-growth ratios. Taken together with previous work and patterns observed in SME, these findings emphasize the role of salt marsh habitat within juvenile blue crab ontogeny and underscore the need to quantify and preserve the complete chain of habitats used by juveniles.

## 1. Introduction

Nursery habitats are critically important for fish and invertebrates. Under the Nursery Role Hypothesis (*sensu* Beck et al., 2001), nursery habitats are characterized by favorable conditions for juveniles, such as higher food availability and lower predation risk, and disproportionately contribute more individuals per unit area to adult segments of the population compared to other habitats (Beck et al., 2001; Heck Jr, Hays and Orth, 2003; Minello et al., 2003; Gillanders et al., 2003). Hence, nursery habitat availability is a major driver of commercially exploited fisheries population dynamics. Consequently, a major research focus in fisheries science and estuarine ecology is identification of nursery habitats for commercially exploited fish and invertebrate species both to prioritize conservation and restoration efforts as well as to guide management decisions (Seitz et al., 2014; Vasconcelos et al., 2014).

Nursery status is often evaluated through four factors: juvenile density, growth, survival, and connectivity between juvenile and adult habitats (Beck et al., 2001). As the contribution of juveniles to adult segments of the population arises from combinations of these four factors, it is generally higher in nursery habitats compared to other candidate nursery habitats (Heck Jr, Hays and Orth, 2003; Minello et al., 2003). However, nursery habitat research typically focuses on one or two of the first three factors due to financial and methodological limitations (Adams et al., 2006; Seitz et al., 2014; Vasconcelos et al., 2014; Litvin et al., 2018; Lipcius et al., 2019).

The Nursery Role Hypothesis maintains that comparisons among all, or at least most, juvenile habitats are required prior to conferring nursery status of a habitat for a given species (Beck et al., 2001; Dahlgren et al., 2006; Litvin et al., 2018). Juveniles tend to utilize structurally complex habitats as nurseries in early life stages in part because of their superior refuge capacity (e.g. Heck Jr, Hays and Orth, 2003; Minello et al., 2003; Lefcheck et al., 2019). The relative value of a given structurally complex habitat as a nursery may be dependent on availability of other habitats with similar characteristics. For example, submersed aquatic vegetation (SAV) or intertidal emergent vegetation (e.g. salt marshes) may seem less important as nurseries in regions where alternative structurally complex habitats are present and accessible (Nagelkerken et al., 2015; Litvin et al., 2018). However, many studies investigating nursery habitats only consider binary comparisons such as a structured habitat and an unstructured control (see review by Ciotti et al., in prep), which can severely limit inference.

Characteristics of nominal nursery habitats (e.g. seagrass meadows or salt marshes) may fluctuate across space and time. The nursery function of these habitats can vary depending on the position within the seascape or season due to the influence of latent environmental, biological, or anthropogenic factors (Nagelkerken et al., 2015; Sheaves et al., 2015; Litvin et al., 2018). For example, predator composition and density vary seasonally in temperate estuaries (Dorenbosch et al., 2009), and may alter habitat use of prey (e.g. Crowder and Cooper, 1982; Fraser and Emmons, 1984; Crowder, Squires and Rice, 1997; Clark, Ruiz and Hines, 2003). Moreover, spatial position within the seascape may modify a habitat’s suitability as a nursery, such as when habitats are positioned close to the site of larval ingress (Stockhausen and Lipcius, 2003) or in areas with low predation pressure (Posey et al., 2005). Assessments of habitats conducted over short temporal intervals or only in one spatial location may miss such phenomena and lead to spurious conclusions about nursery status (Litvin et al., 2018). Hence, these dynamic processes require careful consideration to ensure inferences on habitat comparisons are robust. Moreover, indirect comparisons of multiple habitats via meta-analyses and literature reviews of multiple studies – each considering different combinations of potential nursery habitats – are hindered by the potential influence of confounding, spatiotemporally fluctuating latent variables (e.g. Hyman et al., 2022). Thus, robust evaluation of nursery habitat requires that studies consider as many habitats concomitantly as possible, as well as other influential environmental factors.

Assessment of nursery habitat value is complicated by changes in habitat preferences with ontogeny. Individuals in early-life stages frequently inhabit different habitats than older juveniles or adults (Jones et al., 2010; Nakamura et al., 2012; Epifanio, 2019). Ontogenetic habitat shifts from one nursery to another can minimize mortality-to-growth ratios (*sensu* Werner and Gilliam, 1984), and juvenile survival increases with size (Pile et al., 1996) such that larger juveniles can exploit habitats with less structural refuge and higher food availability (Dahlgren and Eggleston, 2000; Lipcius et al., 2005; Seitz, Lipcius and Seebo, 2005; Nakamura et al., 2012). Consequently, juveniles may utilize different habitats as they grow to minimize mortality-to-growth ratios. Failure to consider these shifts may lead investigators to prioritize only a subset of habitats critical for maintaining healthy population abundances, while neglecting habitats that may be preferred by different stages (Sheaves, Baker and Johnston, 2006; Sheaves et al., 2015; Nagelkerken et al., 2015). Quantitative assessments of nursery function must therefore consider nursery roles within the context of ontogeny, especially for organisms with complex life cycles (Lipcius et al., 2007; Epifanio, 2007; Seitz et al., 2014; Vasconcelos et al., 2014; Litvin et al., 2018; Epifanio, 2019). Partitioning juveniles into multiple size classes and assessing each class concomitantly allows researchers to detect shifts in habitat utilization as juveniles grow and identify stage-specific nursery habitats throughout ontogeny (Nagelkerken et al., 2015; Lugendo et al., 2005; Amorim et al., 2018; Aburto-Oropeza et al., 2009). It is especially important to identify all nurseries used through ontogeny for commercially exploited species with complex life histories so that these habitats can be conserved and related fisheries remain sustainable.

The blue crab *Callinectes sapidus* is an commercially exploited species that relies on structurally complex nursery habitats through ontogeny. The blue crab opportunistically utilizes many habitats in early life stages, including seagrass (e.g. eelgrass *Zostera marina* and widgeon grass *Ruppia maritima* meadows in the Chesapeake Bay), *Spartina alterniflora* salt marshes, and coarse woody debris (see Lipcius et al., 2007, for a review). After re-invading estuaries from the continental shelf, blue crab postlarvae settle into structurally complex nursery habitats, such as seagrass meadows, and rapidly metamorphosize into first instar (j1) juveniles (Epifanio, 2007, 2019). Although some early (j1–j5) juveniles emigrate from initial settlement locations to avoid adverse density-dependent effects associated with conspecifics (Etherington and Eggleston, 2000; Blackmon and Eggleston, 2001; Reyns and Eggleston, 2004), many remain to exploit the high refuge quality afforded by primary nursery grounds. As juveniles outgrow the mouth-gape sizes of smaller predators, they emigrate to other habitats with lower quality refuge but more abundant preferred prey (e.g. *Macoma balthica*; Seitz et al., 2003; Lipcius et al., 2005; Seitz, Lipcius and Seebo, 2005).

Several studies have posited different size thresholds for when emigration out of primary nursery grounds unfolds. A mesocosm experiment examining the effects of simulated *S. alternaflora* shoots on survival estimated that juvenile blue crabs may shift their habitat preferences at sizes as small as 12 mm carapace width (CW), when they could achieve a size refuge from smaller predators abundant within salt marsh habitats (e.g. *Fundulus heteroclitus* Orth and van Montfrans, 2002). Subsequent field studies maintained that juveniles begin emigrating from seagrass meadows to utilize unstructured and salt marsh habitats only after reaching 25–30 mm CW (e.g. Pile et al., 1996; Lipcius et al., 2005; Johnston and Lipcius, 2012; Hyman et al., 2022). Notably, these hypotheses are not mutually exclusive. Salt marsh habitat may represent an intermediate nursery – one with marginally lower refuge quality than seagrass but higher food availability (Seitz et al., 2003; Seitz, Lipcius and Seebo, 2005) – before juveniles emigrate to unstructured or alternative nursery habitats (e.g. Stockhausen and Lipcius, 2003; Mizerek, Regan and Hovel, 2011; Ralph, 2014; Wood and Lipcius, 2022).

Secondary production from a habitat is the common currency used to quantify the value of different habitats (Peterson and Lipcius, 2003). Small-scale studies previously demonstrated that blue crab production from salt marshes was substantial in the Gulf of Mexico (Thomas, Zimmerman and Minello, 1990a; Zimmerman, Minello and Rozas, 2000) and Chesapeake Bay (Cicchetti, 1998; Cicchetti and Diaz, 2000). Recently, large-scale spatiotemporal analyses of juvenile blue crab habitats emphasized the roles of salt marsh and high-turbidity habitats in addition to seagrass meadows in promoting secondary production (Hyman et al., 2022). Spatially explicit analyses of relative secondary production are useful for assessing potential nursery capacity of large regions (Forrester and Swearer, 2002; Berkström et al., 2013; Ralph et al., 2013). For example, by exploiting ontogenetic shifts in habitat usage, such as juveniles emigrating from nursery habitats to unstructured adult habitats, broad-scale studies can highlight productive areas which can be prioritized for conservation (e.g. Hyman et al., 2022). However, evaluation of nursery habitats at broad scales may miss important processes operating at smaller scales. In addition, broad-scale studies are unable to ascertain which aspects of a habitat promote secondary production. For example, although turbid salt marsh habitat is positively associated with juvenile blue crab density at large spatial and temporal scales (Hyman et al., 2022), it is unclear if such production is more closely linked to vegetative structure (i.e. *Spartina* shoots; Johnson and Eggleston, 2010; Isdell et al., 2021), or to structurally complex detritus along erosional marsh shorelines (Etherington and Eggleston, 2000; Etherington, Eggleston and Stockhausen, 2003). As small-scale studies are uniquely suited for capturing these processes, it is important to employ this approach in concert with broad-scale studies to determine: (1) which habitats are associated with high juvenile density; (2) which habitat characteristics are important in promoting juvenile density, and (3) which environmental variables modify habitat suitability.

In this study, we modeled juvenile blue crab abundance of two size classes (herein small: *≤*15 mm CW, and large: 16–30 mm CW) in multiple juvenile habitats at various locations within an estuarine seascape during the blue crab late summer-fall recruitment season. Building on previous work (Hyman et al., 2022), we sought to complement large-scale spatiotemporal analyses with mensurative experiments (*sensu* Underwood, Chapman and Connell, 2000) at local spatial (i.e. 10s of kilometers) and temporal (i.e. biweekly) scales within the York River, a tributary of Chesapeake Bay. Specifically, we wanted to determine the effects of habitat, spatial position, and environmental factors on juvenile blue crab density. To accomplish this, we developed multiple models (labeled *g_i_*) with different combinations of spatial position, habitat, and turbidity as independent variables. We describe and justify the models and corresponding independent variables in Appendix A.

## 2. Habitats considered

We examined four habitat types—unstructured sand, seagrass meadows, salt marsh edge, and shallow detrital habitat. Seagrass meadows (herein, seagrass) are regarded as preferred nursery for juvenile blue crabs (e.g. Orth and van Montfrans, 1987; Perkins-Visser, Wolcott and Wolcott, 1996; Hovel and Lipcius, 2002; Hovel and Fonseca, 2005; Ralph et al., 2013) due to disproportionately high densities and survival of small juvenile crabs (i.e. *<*30 mm CW) in seagrass meadows relative to other potential nursery habitats (e.g. Orth and van Montfrans, 1987; Pile et al., 1996; Lipcius et al., 2005). Meanwhile, salt marshes serve as alternative nursery habitat for juvenile blue crabs in locations where seagrass is absent or declining (Fitz and Wiegert, 1991; Jivoff and Able, 2003; Bishop et al., 2010; Johnson and Eggleston, 2010). In Gulf of Mexico and some Chesapeake Bay nursery habitats, juvenile blue crab density was high in both seagrass and salt marsh habitats (Thomas, Zimmerman and Minello, 1990b; Rozas and Minello, 1998; Heck Jr, Coen and Morgan, 2001; Hyman et al., 2022). Salt marshes may afford refuge through structurally complex shoots and rhizomes (i.e. salt marsh edge; SME; Johnson and Eggleston, 2010; Miller et al., 2023). In addition, detritus exported from the vegetated marsh surface accumulates in adjacent tidal marsh creeks. This “shallow detrital habitat” (SDH) is associated with eroding peat and can harbor high densities of juvenile blue crabs (Etherington and Eggleston, 2000; Etherington, Eggleston and Stockhausen, 2003; Voigt and Eggleston, 2022). Finally, unstructured sand habitat (herein, sand) constitutes most available shallow habitat in the Chesapeake Bay, but is characterized by relatively low predation refuge (Lipcius et al., 2005) and serves as a control to assess nursery value of seagrass and salt marsh habitats for juvenile blue crabs (e.g. Heck Jr, Coen and Morgan, 2001; Lipcius et al., 2005; Shakeri et al., 2020).

## 3. Methods

### 3.1. Study Area

Field work was conducted in the York River, a tributary in the lower portion of western Chesapeake Bay between August and November, 2020. The river is morphometrically characterized by depths generally between 5 to 10 m along the axes, but with deeper portions (>20 m) near the mouth (Smock, Wright and Benke, 2005). In addition, This system contains a range of seagrass, salt marsh, and unstructured sand habitat configurations ideally suited for investigating the relative importance of multiple habitat types (Hovel and Lipcius, 2002; Lipcius et al., 2005). Seagrasses, primarily eelgrass (*Zostera marina*) and Widgeon grass (*Ruppia maritima*), vary from large, continuous meadows to areas with few small patches of variable shoot densities (Hovel and Lipcius, 2002). Salt marshes, dominated by smooth cordgrass (*Spartina alterniflora*), span extensive sections of the shorelines, although areal coverage of marsh patches varies spatially along the shorelines. Secchi disk depth values, a proxy for turbidity, range from 0.5–1.5m at the mouth of the system and 0–0.5 m upriver near the confluence of the Pamunkey and Mattaponi tributaries. For a more detailed description of physiochemical variables, see Table S1. The river was divided into 3 approximately evenly split strata (nearly 17 km each) for a lack of an obvious stratification strategy, constituting downriver, midriver, and upriver strata (Fig. 1).

**FIG 1.**
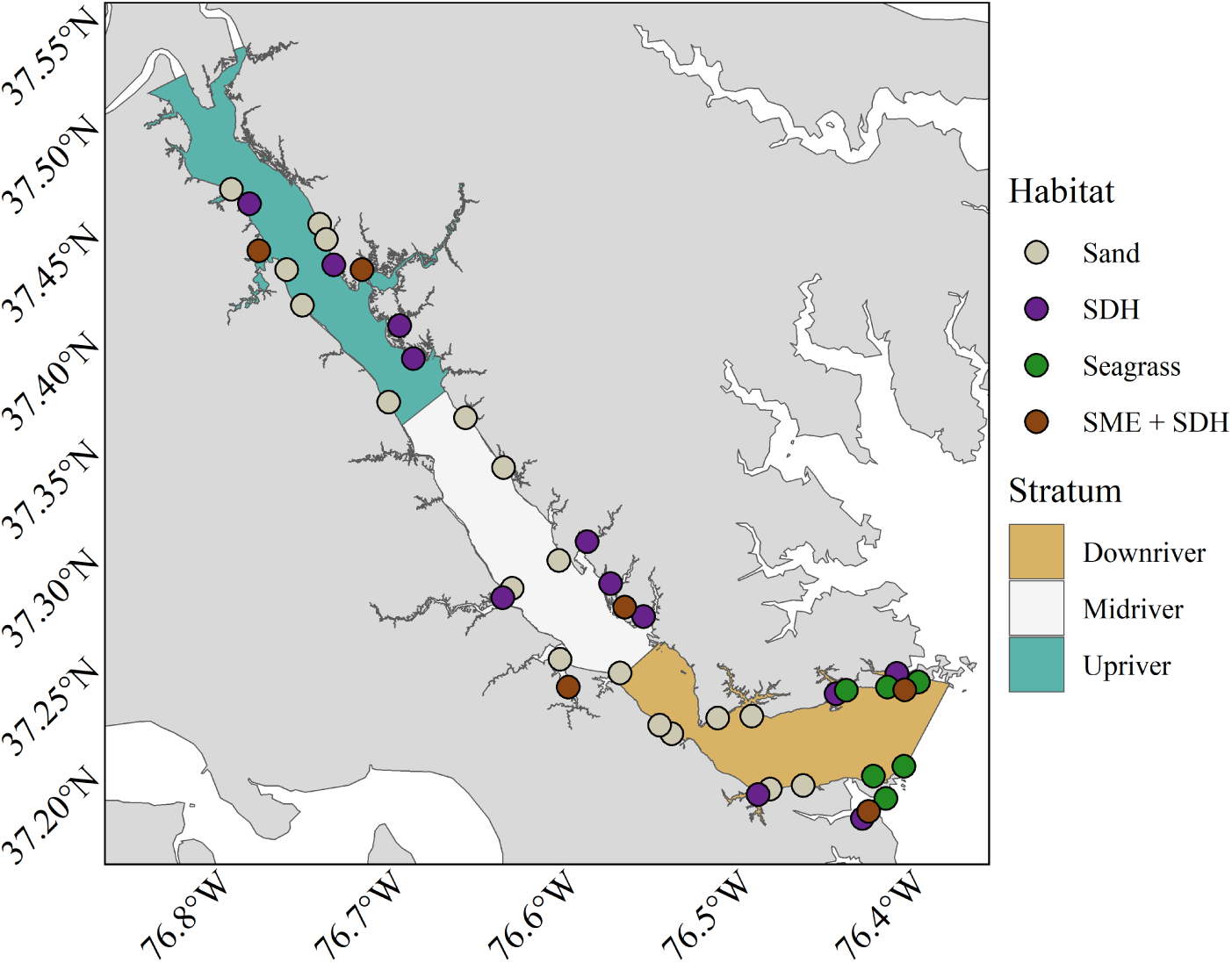
Map of the York River displaying sampling sites colored by habitat type. SME sites are a subset of SDH sites.

### 3.2. Sampling design

Site selection was achieved via a random sampling algorithm. Selection involved (1) extracting geographic coordinates for the entire shoreline of the York River, (2) subsetting coordinates by habitat type and stratum, and (3) randomly selecting a prespecified number of stations for each habitat within each stratum. Six SDH and sand stations were selected in each stratum, while two SME stations were randomly selected from the six SDH sites in each stratum. Finally, six seagrass stations were randomly selected from the downriver stratum, as seagrass is absent in midriver or upriver strata (Fig. 1). The number of sites per habitat in each stratum were the maximum logistically feasible to sample in a day given time constraints and tidal considerations.

Between August and November 2020, juvenile blue crabs were sampled in seagrass, SME, SDH, and sand at biweekly intervals. Four sampling trips were conducted to sample all four habitats. The first three sampling trips – targeting seagrass, SDH, and sand – were conducted August 24^th^–27^th^ (Trip 1), September 15^th^–22^nd^ (Trip 2), and October 5^th^–8^th^ (Trip 3). SME was also sampled on trips 2 and 3, as well as Trip 4, which occurred October 19^th^–20^rd^. Hence, there is confounding between trip 4 and SME habitat, otherwise exploratory data analyses did not indicate interactions between habitat and trip. This culminated in a total of 144 samples (Table S2), although five samples were later expunged due to missing predictor values (i.e. Secchi disk depth) in seagrass (two) and SDH (three).

Each habitat was sampled using gear and methodologies corresponding to habitat-specific structure and bottom types. All gear types used 3-mm mesh netting to ensure that size-specific catchability was consistent after accounting for differences in gear efficiency. SDH and sand stations were sampled ± 3 h of high tide via benthic scrapes towed for 20 m along tidal salt marsh creek and beach shorelines, respectively (see Ralph and Lipcius, 2014, for details). Meanwhile, SME stations were sampled using modified flume nets set at flood tide and collected at ebb tide (Fig. S1; McIvor and Odum, 1986). At seagrass stations, a 1.68-m^2^ drop-cylinder and a 10-cm diam PVC suction pipe attached to a sampling pump, modified from Orth and van Montfrans (1987), were employed to collect juvenile blue crabs (e.g., Orth and van Montfrans, 1987; Ralph et al., 2013; Hovel and Lipcius, 2002; Heck Jr, Coen and Morgan, 2001). Seagrass stations were suctioned within the drop-cylinders for 6 min continuously. For sand, SDH, and SME stations, immature juvenile crabs were counted, measured *in situ* and released. The contents of each seagrass suction sample were frozen for storage and subsequently examined for juvenile blue crabs, double-checked, and all crabs counted and measured. Physicochemical variables salinity, temperature, and turbidity were recorded using a YSI data sonde (for salinity and temperature) and a Secchi disk (water clarity, the inverse of turbidity) at each station on each trip.

### 3.3. Analyses

#### 3.3.1. Basic model structure

All data analyses, transformations, and visualizations were carried out using the R programming language for statistical computing (R Core Team, 2022). Relationships between both small and large juvenile blue crab abundance and environmental variables were evaluated using multivariate negative binomial linear mixed-effects models within a Bayesian framework. The predictor variables for juvenile abundance include habitat (seagrass, SME, SDH, and sand), spatial stratum (downriver, midriver, and upriver), and turbidity.

Transformations to turbidity values were applied prior to their inclusion in abundance models. Here, ln turbidity was defined as the natural log transformation of Secchi-disk depth, multiplied by -1 (*T* = *−* ln Secchi). The natural log transformation was applied based on the assumption that a threshold exists in water transparency. Assuming that effects of turbidity on juvenile abundance reflect refuge from visually oriented predators (top-down control), small changes in water transparency when water is relatively clear are not expected to substantially affect juvenile abundance as much as small changes in water transparency when water is turbid (e.g. predation rates by summer flounder on mysid shrimp; Howson, 2000). Similarly, if associations between juvenile abundance and turbidity are related to elevated food availability near the estuarine turbidity maximum, juveniles would presumably remain more sensitive to fluctuations in turbidity at high values compared to clearer waters. Multiplying the variable by -1 facilitates inference on turbidity, instead of water transparency (inverse).

For the *s*^th^ site on trip *t* in habitat *h*, the Bayesian model for juvenile blue crab abundance of size class *i* is expressed as:

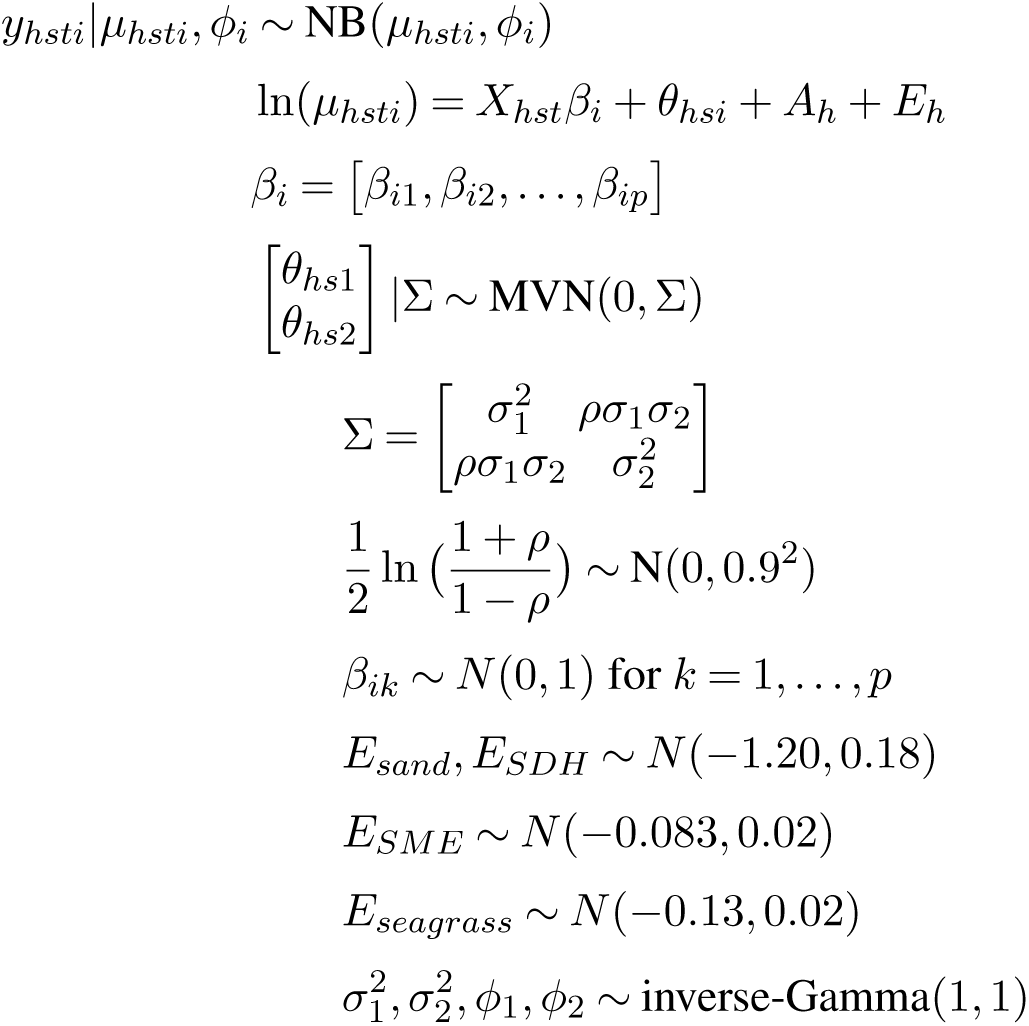

where *NB*(*µ_hsti_, ϕ_i_*) denotes a negative binomial type II distribution with mean *µ_hsti_*, while *ϕ_i_* controls the over-dispersion for each size class such that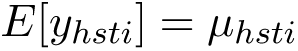 and 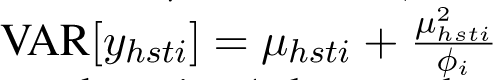. The response variables, juvenile crab counts for size classes, are denoted *y_hsti_* where *i* = 1 denotes the small size class (*≤* 15 mm) and *i* = 2 denotes the large size class (16–30 mm). Total area sampled (seagrass = 1.68 m^2^, SME = 1 m^2^, SDH and sand = 20 m^2^) is included as an offset term *A_h_*. Here, *θ_hs_* denotes a site-specific random effect following a multivariate normal (*MV N*) distribution with mean 0 and co-variance Σ. The covariance matrix is composed of 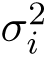 along the diagonal denoting variance for size class *i* and *ρ* describing the correlation between size classes at a given site. A normal prior was applied to Fisher-transformed *ρ* to constrain values between -1 and 1. In addition, due to varying requirements as a function of size, each size class was not expected to respond equally to predictor variables *X_hst_* (habitat, turbidity, spatial position, and relevant interaction terms, see Appendix A for details). Hence, *β_i_* refer to regression coefficients for each size class *i* associated with *X_hst_*. Measurements of both the abundances of size classes and predictors predictors *X_hst_* were taken at the site-trip spatiotemporal resolution, such that predictors were not specific to any one size class *i* but to all sizes classes at a given site-trip. Informative prior distributions for gear efficiency *E_h_* were supplied based on gear efficiencies from literature (seagrass, SDH, and sand) and from the fall pilot study (SME), converted to the ln-scale (see Appendix B for details). This incorporated increased uncertainty into habitat-specific estimates.

Bayesian inference required numerical approximation of the joint posterior distribution of all model parameters including the vectors of random effects. To this end, we implemented the model using the Stan programming language for Bayesian inference to generate Markov chain Monte Carlo (MCMC) samples from the posterior (Gelman, Lee and Guo, 2015). For each model, we ran four parallel Markov chains, each with 5,000 iterations for the warm-up/adaptive phase, and another 5,000 iterations as posterior samples (i.e. 20,000 draws in total for posterior inference). Convergence of the chains was determined both by visual inspection of trace plots (e.g. Fig S2) and through inspection of the split *R*^^^ statistic. All sampled parameters had an *R*^^^ value less than 1.01, indicating chain convergence (Gelman, Lee and Guo, 2015). We considered covariates and interactions whose posterior distributions indicated a positive or negative effect with *≥* 80% posterior probability, as scientifically relevant to juvenile blue crab abundance (Kruschke, 2021). All CIs referenced here are the highest posterior density intervals (McElreath, 2018).

Estimated log-pointwise predictive density (ELPD) and related Δ*_ELP D_* values were used to evaluate the degree of predictive power for each model among the set of statistical models *g_i_* (Vehtari, Gelman and Gabry, 2017). Values of ELPD are widely employed to measure out-of sample predictive accuracy, while Δ*_ELP D_* values refer to the difference in ELPD between a given model and the model with the best ELPD in the set. Values of ELPD and Δ*_ELP D_* were estimated using the Widely-Applicable Information Criterion (WAIC; Watanabe, 2013; Gelman, Lee and Guo, 2015; Vehtari, Gelman and Gabry, 2017). Both WAIC and ELPD were estimated using the `loo` package (Vehtari et al., 2022). When two models had comparable Δ*_ELP D_* values (i.e. *≤* 4; Sivula et al., 2020), the simpler model was chosen as the more appropriate model under the principle of parsimony.

#### 3.3.2. Alternative model structures

In our design, the downriver stratum was partially confounded with seagrass habitat, as seagrass is present only at the mouth of the York River (Fig. 1; Hyman et al., 2022). As a consequence, under this design it is not easily discernible whether spatial stratum interacted with seagrass habitat. To ensure that seagrass habitat and spatial stratum did not interact and influence results, we constructed two additional models – one with only SME, SDH, and sand across all strata and a second with all four habitats only in the downriver stratum – and compared the results to our best-fitting model, *g*_1_. Posterior distributions of main effects (where present across models) strongly overlapped, indicating that interactions between spatial stratum and habitat were unlikely when other predictors were considered (Fig. S3).

#### 3.3.3. Conditional effects

Conditional effects plots were used to visualize the relationship between response variables (juvenile blue crab abundance) and meaningful predictors both among habitats within a size class and within habitats between size classes. Herein, we refer to “conditional effects“ as the effects of a given predictor (either continuous or categorical) while holding all random effects at 0 and fixing co-varying predictors. Specifically, we held ln turbidity at 0 to estimate habitat conditional effects and held habitat effects at the reference (i.e. sand; h = 1). Conditional effects were used to conceptualize mean effects of each level in a given categorical variable. Hence conditional linear contrast statements were used to determine whether differences in abundances among habitats were statistically meaningful. For the *h*^th^ habitat (where *h >* 1), we considered pairwise difference between habitats *β_hi_ − β*_1_*_i_*, where 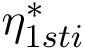 is the reference intercept (sand). Meanwhile, for comparisons of within-habitat abundances between *≤* 15 (i.e. *i* = 15) and 16–30 mm size classes, for the *h^th^* habitat, we considered the contrast *β_h,_*_15_ *−β_h,_*_30_.

## 4. Results

We collected and measured 1,004 juvenile blue crabs *≤* 30 mm CW from 139 samples. A complete summary of all physicochemical variables and crab sizes is detailed in Table S1, while a histogram of crab sizes is provided in Figure S4. Herein, all abundance values for size classes refer to abundance per square meter, and are referred to as density.

### 4.1. Model selection

The best fitting model was *g*_5_, which posited juvenile blue crab abundance as a function of habitat, turbidity, stratum, and a habitat-turbidity interaction. However, all models had comparable ELPD values (Δ*_ELP D_ ≤* 4 for all models except *g*_4_) and overlapping standard errors (Table 1), indicating relative statistical equivalence. Hence, we chose the simplest model *g*_1_, with habitat and turbidity additive, as the best model under the principle of parsimony. Hereafter, inferences are based on model *g*_1_.

**TABLE 1.** *Model selection results from five Bayesian multivariate negative binomial regression models (g_i_) using* ln *turbidity (**T**), habitat (**H**), and stratum (**S**) as predictors of juvenile blue crab abundance. Models are presented in order of predictive power based on collected data. **WAIC**: the Widely-Applicable Information Criterion; **ELPD_WAIC_**: the estimated log-pointwise density calculated from WAIC;* Δ***_ELPD_****: the relative difference between the ELPD of any model and the best model in the set; SE*_Δ*ELPD*_ *: standard error for the pairwise differences in ELPD between the best model and any given model; **pWAIC**: estimated effective number of parameters. The selected model (g*_1_*) values are presented in bold font. Model justifications are in Appendix A*

| Model: Fixed effects in mean structure | WAIC | ELPD<sub>WAIC</sub> | ΔELPD | $SE_{\Delta ELPD}$ | pWAIC |
| --- | --- | --- | --- | --- | --- |
| $g_5$: H + T + S + (H x T) | 1049.76 | -524.88 | 0.00 | 0.00 | 37.03 |
| $g_1$: **H + T** | **1050.46** | **-525.23** | **-0.35** | **3.31** | **36.92** |
| $g_2$: H + T + (H x T) | 1050.60 | -525.30 | -0.42 | 1.67 | 37.36 |
| $g_3$: H + T + S | 1050.76 | -525.38 | -0.50 | 3.17 | 37.13 |
| $g_4$: H + T + S + (H x S) | 1058.09 | -529.04 | -4.16 | 3.71 | 40.43 |

### 4.2. Habitat effects

#### 4.2.1. Small (≤15 mm) size class

Small juvenile blue crab density was highest in seagrass (13.83 per m^2^ on the count scale), followed by SME (4.70), SDH (0.62), and sand (0.35) (Table 2; Figs. 2 and S5). For pairwise linear contrasts among habitats, the posterior probability that a given contrast was positive all exceeded 90%, indicating that differences in the expected density of small juvenile crabs among habitats were statistically meaningful (Table 3 and Fig. S6).

**TABLE 2.** *Posterior summary statistics (median and 80% CI) of habitat and turbidity effects for the small (≤15 mm CW) juvenile size class based on model g*_1_*. Habitat values represent the expected small juvenile density in a given habitat (abundance per m*^2^*). Meanwhile, the last column reflects the effect (i.e. regression coefficient) of* ln *turbidity on small juvenile density, irrespective of habitat. Values are supplied on both the model (*ln*) and count scales*.

| Scale | Quantile | Sand | Seagrass | SME | SDH | $\ln$ Turbidity |
| --- | --- | --- | --- | --- | --- | --- |
| Model | 10% | -2.02 | 1.24 | 0.38 | -1.54 | -0.06 |
|  | 50% | -1.06 | 2.63 | 1.55 | -0.48 | 0.18 |
|  | 90% | -0.10 | 3.94 | 2.66 | 0.58 | 0.43 |
| Count | 10% | 0.13 | 3.47 | 1.47 | 0.21 | 0.94 |
|  | 50% | 0.35 | 13.83 | 4.70 | 0.62 | 1.20 |
|  | 90% | 0.91 | 51.39 | 14.30 | 1.78 | 1.54 |

**FIG 2.**
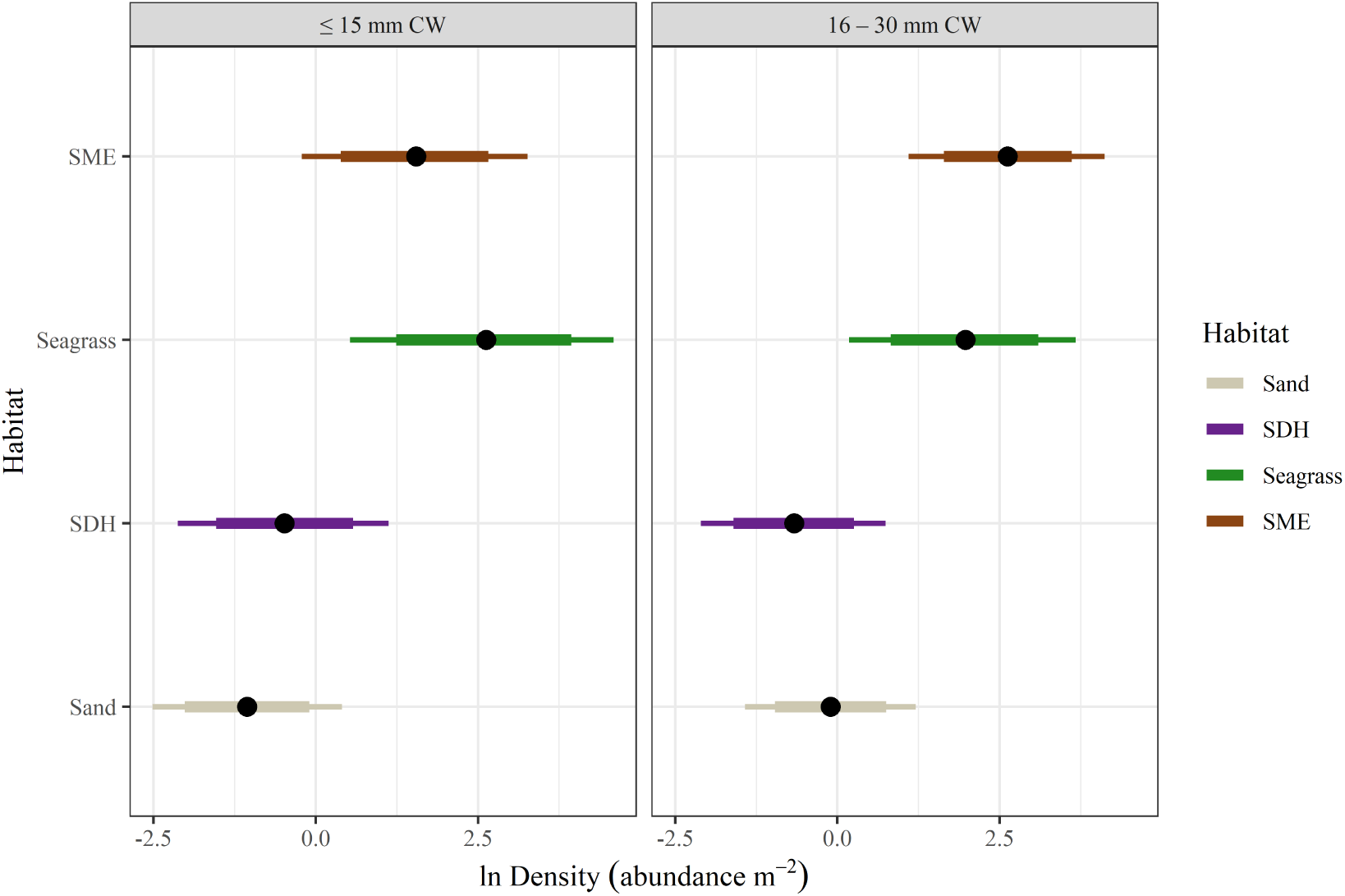
Posterior distributions of habitat-specific conditional ln expected densities (holding random effects and ln turbdity at 0), from model g_1_ for both small (≤15 mm CW; left column) and large (16–30 mm CW; right column) size classes. Dots denote posterior median expected values, while thick bars represent 80% Bayesian CI’s and thin bars denote 95% Bayesian CI’s.

**TABLE 3.** Within-size class linear contrast depicting differences in expected juvenile blue crab density between habitats from Model g_1_ (see Section 3.3.3). Percentages indicate 80% CI and median of differences in effect sizes, while the final two columns list the probability of a positive or negative effect.

| Size Class | Contrast | 10% | 50% | 90% | Pr > 0 | Pr < 0 |
| --- | --- | --- | --- | --- | --- | --- |
| Small | SDH – Sand | 0.05 | 0.59 | 1.09 | 0.92 | 0.08 |
| | Seagrass – Sand | 2.83 | 3.70 | 4.47 | $\approx 1.00$ | $\approx 0.00$ |
| | Seagrass – SDH | 2.33 | 3.11 | 3.83 | $\approx 1.00$ | $\approx 0.00$ |
|  | Seagrass – SME | 0.28 | 1.10 | 1.85 | 0.95 | 0.05 |
| | SME – Sand | 1.94 | 2.60 | 3.24 | $\approx 1.00$ | $\approx 0.00$ |
| | SME – SDH | 1.52 | 2.01 | 2.51 | $\approx 1.00$ | $\approx 0.00$ |
| Large | SDH – Sand | -0.97 | -0.56 | -0.17 | 0.03 | 0.97 |
| | Seagrass – Sand | 1.44 | 2.08 | 2.68 | $\approx 1.00$ | $\approx 0.00$ |
| | Seagrass – SDH | 2.02 | 2.64 | 3.23 | $\approx 1.00$ | $\approx 0.00$ |
|  | Seagrass – SME | -1.27 | -0.65 | -0.07 | 0.07 | 0.93 |
| | SME – Sand | 2.21 | 2.73 | 3.24 | $\approx 1.00$ | $\approx 0.00$ |
| | SME – SDH | 2.84 | 3.29 | 3.76 | $\approx 1.00$ | $\approx 0.00$ |

#### 4.2.2. Large (16-30 mm) size class

In contrast to the smaller size class, density of the large size class was highest in SME (13.80), followed by seagrass (7.19), sand (0.90), and SDH (0.51) (Table 4; Fig. 2). For pairwise linear contrasts among habitats SME–SDH, SME–sand, seagrass–SDH, and seagrass–sand, the posterior probability that a given contrast was positive exceeded 90%. Meanwhile, for pairwise linear contrasts among habitats seagrass–SME and SDH–sand, the posterior probability that a given contrast was negative exceeded 90%. Taken together, these results indicated that differences in the expected density of small juvenile crabs among habitats were statistically meaningful (Table 3 and Fig. S6).

**TABLE 4.** *Posterior summary statistics (median and 80% CI) of habitat and turbidity effects for the large (16–30 mm CW) juvenile size class based on model g*_1_*. Habitat values represent the conditional expected large juvenile density in a given habitat (see Section 3.3.3). Meanwhile, the last column reflects the effect (i.e. regression coefficient) of* ln *turbidity on large juvenile density, irrespective of habitat. Values are supplied on both the model (*ln*) and count scales*.

| Scale | Quantile | Sand | Seagrass | SME | SDH | $\ln$ Turbidity |
| --- | --- | --- | --- | --- | --- | --- |
| Model | 10% | -0.96 | 0.82 | 1.64 | -1.60 | 0.07 |
|  | 50% | -0.11 | 1.97 | 2.62 | -0.67 | 0.29 |
|  | 90% | 0.75 | 3.10 | 3.61 | 0.26 | 0.51 |
| Count | 10% | 0.38 | 2.27 | 5.16 | 0.20 | 1.07 |
|  | 50% | 0.90 | 7.19 | 13.80 | 0.51 | 1.33 |
|  | 90% | 2.12 | 22.09 | 37.00 | 1.29 | 1.67 |

Pairwise linear contrasts among habitats yielded posterior probabilities that a given contrast was pos-itive or negative all exceeded 90%, indicating that differences in the expected density of large juvenile crabs among habitats were statistically meaningful (Table 3 and Fig. S7).

#### 4.2.3. Comparisons among size classes

Within-habitat linear contrasts between small and large size classes indicated changes in habitat utilization with size. Moving from small to large size classes, utilization decreased in seagrass meadows, increased in both SME and sand, and did not change appreciably in SDH. The probability of seagrass harboring fewer large crabs than small crabs was 70%, indicating weak-moderate support but failing to meet our threshold for relevance. Meanwhile, the probability that SME and sand harbored more large individuals than small individuals were both 85% (Table 5; Fig. S8). Conversely, contrasts among size classes for SDH were distributed evenly across both negative and positive values, indicating considerable uncertainty and no discernible size effect.

**TABLE 5.** Within-habitat linear contrasts depicting differences in expected juvenile blue crab density for small and large size class (see Section 3.3.3) Positive values indicate increases in expected density as animals grow from ≤15 to 16 – 30 mm, while negative values indicate decreases in expected density. The first three rows indicate 80% CI and median values, while the final two rows list the probability of a positive or negative effect.

| Quantile | Sand | Seagrass | SME | SDH |
| --- | --- | --- | --- | --- |
| 10% | -0.24 | -2.22 | -0.26 | -1.46 |
| 50% | 0.94 | -0.66 | 1.07 | -0.21 |
| 90% | 2.17 | 0.95 | 2.47 | 1.12 |
| Probability |  |  |  |  |
| Pr > 0 | 0.85 | 0.30 | 0.85 | 0.42 |
| Pr < 0 | 0.15 | 0.70 | 0.15 | 0.58 |

### 4.3. Turbidity effects

Turbidity was positively associated with juvenile density, though the effect of turbidity was stronger for large juveniles. Posterior distributions of regression coefficients for ln turbidity indicated a broadly positive effect for both small and large size classes. The probability that the effect sizes of turbidity were positive were 83% and 96% for small and large juveniles, respectively (Tables 2 and 4; Fig. 3 right of red line).

**FIG 3.**
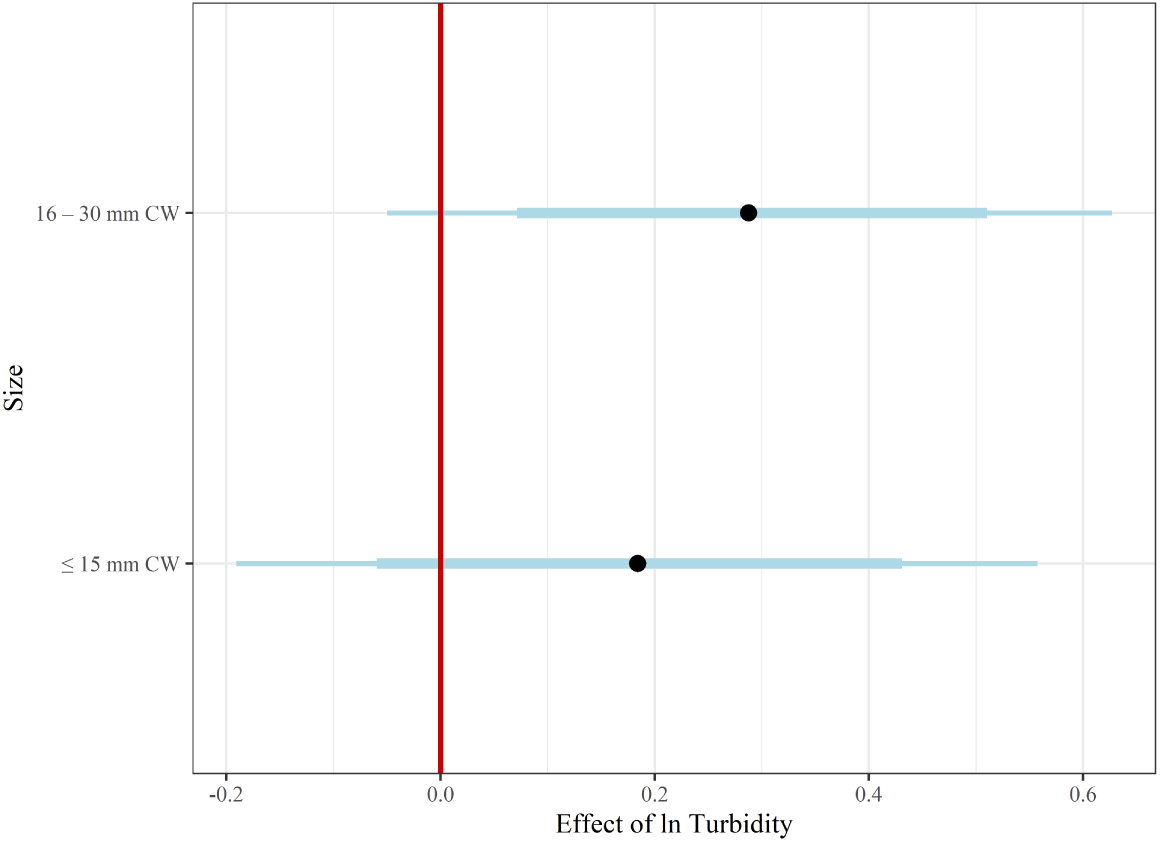
Posterior summaries (median and CIs) for ln turbidity regression coefficients for small (≤15 mm CW) and large (16–30 mm CW) size classes. Dots denote posterior median difference in expected values, while thick bars represent 80% Bayesian CI’s and thin bars denote 95% Bayesian CI’s. The red line denotes 0.

### 4.4. Correlation between size classes

The posterior distribution of *ρ* suggested that substantial dependence existed among size classes (Fig. 4). The posterior distribution of *ρ* yielded median (80% CI) of 0.90 (0.71–0.97), which indicated strong positive associations between size classes.

**FIG 4.**
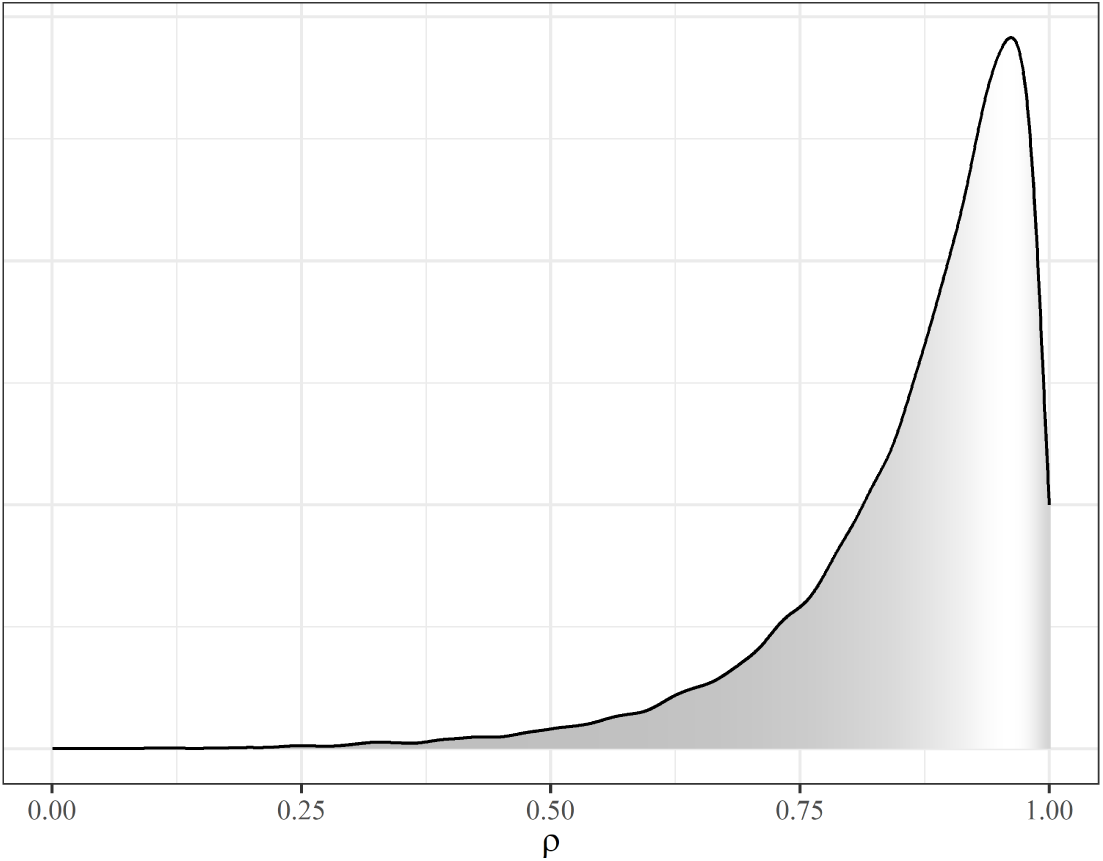
Posterior distribution for the correlation parameter ρ between small and large juvenile size classes.

## 5. Discussion

This study presents model-based evidence of differential habitat utilization among juvenile blue crab early-life stages and suggests the current paradigm surrounding blue crab early life history requires revision. Subsetting young-of-year juveniles into finer-scale size classes enabled us to observe shifts in density between small and large size classes among habitats, particularly from seagrass to SME. Our results are consistent with previous work emphasizing seagrass as an important nursery for the smallest juveniles (e.g., Lipcius et al., 2007), and suggest that SME represents a possible intermediate nursery habitat following initial emigration from seagrass beds but before occupying unstructured habitat commonly utilized by adults (Lipcius et al., 2005). After accounting for habitat-specific differences in density, turbidity was positively related with both small and large juveniles. However, our models could not address whether this effect is due to top-down (predation) or bottom-up (food availability) controls. Although more evidence (i.e., comparisons of survival and growth) is required to ascertain the exact role of structured marsh habitat in juvenile blue crab ecology, we posit that both seagrass and SME habitats are important in maintaining adult populations and serve as nurseries for different size classes of juveniles.

### 5.1. Size-specific habitat effects

Small juvenile blue crab density was highest in structurally complex seagrass and SME habitats. In our study, seagrass meadows harbored the highest densities of small juvenile crabs, which is consistent with previous work emphasizing this habitat as the preferred nursery for small juveniles (Orth and van Montfrans, 1987; Perkins-Visser, Wolcott and Wolcott, 1996; Hovel and Lipcius, 2002; Ralph et al., 2013; Voigt and Eggleston, 2022). We also observed that SME harbored high densities of small juveniles per square meter relative to sand and SDH, although SME densities remained much lower than those estimated in seagrass. Postlarvae re-invading Chesapeake Bay likely encounter seagrass beds and other SAV such as the non-native macroalga *Gracilaria vermiculophylla* and preferentially choose these habitats for initial settlement (Stockhausen and Lipcius, 2003; van Montfrans, Ryer and Orth, 2003; Johnston and Lipcius, 2012; Wood and Lipcius, 2022). However, heterogeneity in hydrodynamic conditions can cause a substantial proportion of ingressing postlarvae to miss structurally complex SAV habitats (Stockhausen and Lipcius, 2003). Although somewhat less suitable than SAV, SME provides an alternative nursery habitat. In addition, a proportion of early juveniles in SAV emigrate to alternative substrates to avoid adverse density-dependent effects (Etherington and Eggleston, 2000; Reyns and Eggleston, 2004; Blackmon and Eggleston, 2001). High densities of juveniles observed in salt marshes at all spatial locations within the tributary likely reflect a combination of these two processes.

In contrast, small juvenile densities in SDH and sand were less than 1 m*^−^*^2^, suggesting that these habitats were relatively unproductive. Although food availability can be high in sand, occupation by smaller juveniles in this habitat is likely discouraged by low structural refuge. High densities of small juveniles were reported inhabiting SDH in North Carolina estuaries (e.g. Etherington and Eggleston, 2000; Etherington, Eggleston and Stockhausen, 2003; Voigt and Eggleston, 2022). We estimated far lower small juvenile densities in similar habitat in the York River. It is unclear why SDH is an attractive habitat for small juveniles in other locations but not within the York River, although differences in gear type or hydrodynamics associated with wind-driven vs tidally-driven estuaries may be responsible for these discrepancies. Specifically, logistical issues related to benthic scrapes may make this gear type inefficient when assessing abundance in SDH. Unlike sand, SDH is characterized by pitted surfaces and complex material, such that this gear type may be much less efficient in this habitat than efficiency estimates would indicate. As a result, we caution that our abundance estimates may be overly conservative, and stress that additional studies using gear better suited to sample SDH (Voigt and Eggleston, 2022) are required to validate our estimates.

Our estimated habitat-specific abundance patterns changed notably between small and large juveniles. Whereas small juveniles were more abundant in seagrass meadows than in SME, this pattern reversed among large juveniles. Moreover, large juveniles were more abundant in SME and less abundant in sea-grass relative to small juveniles. Decreases in habitat-specific density of large juveniles with size are understood as being due to mortality or emigration of small juveniles. Mortality and emigration may explain decreases in juvenile abundance between small and large size classes, but the degree to which these processes affect density is not clear. In contrast, increases in the density of large juveniles in SME are indicative of a possible shift in habitat utilization concurrent with losses due to mortality and emigration, and consequently the preference of SME for this size class may be understated.

Patterns in size-specific habitat utilization observed here likely result from changing requirements of juvenile blue crabs with size. Seagrass meadows afford high survival to newly settled juveniles, particularly from smaller predators, due to the small interstitial spaces between shoots and rhizomes. In contrast, emergent salt marsh vegetation has higher interstitial space between shoots, allowing small predators such as the mummichog (*Fundulus heteroclitus*) to navigate and forage within the inundated marsh surface (Orth and van Montfrans, 2002). In the absence of juvenile density-dependent effects, smaller juveniles may prefer seagrass meadows because of the lower mortality risk when compared to salt marshes. However, upon reaching 10–15 mm CW, juvenile blue crabs outgrow the mouth-gape sizes of many smaller predators, making salt marsh habitats favorable (Orth and van Montfrans, 2002; Urban, 2007). Furthermore, marsh shoots are dense enough to prevent larger predators from foraging effectively (Johnston and Caretti, 2017; Miller et al., 2023). Salt marshes additionally harbor abundant detrital material, bivalves, and other invertebrates, which are consumed by juveniles to accelerate growth (Seitz, Lipcius and Seebo, 2005). The combination of lower mortality risk from small predators and high food availability is consistent with mechanisms driving ontogenetic shifts in many marine species (Werner and Gilliam, 1984; Dahlgren and Eggleston, 2000), and accounts for shifts in utilization from seagrass to SME as juveniles grow.

Similar to patterns we observed in SME, sand utilization also increased among large juveniles. Estimated densities of large juveniles in sand were nearly triple those of smaller juveniles, although in both size classes, abundance in sand was much lower than in seagrass and SME. These findings are consistent with the current paradigm that as juveniles reach 20–30 mm CW, they reach a size refuge from a broader suite of predators and are free to increasingly exploit unstructured habitat with less refuge but high food availability (Lipcius et al., 2005; Seitz, Lipcius and Seebo, 2005).

Taken together with previous work and patterns observed in SME (Lipcius et al., 2005; Johnson and Eggleston, 2010; Hyman et al., 2022), our findings suggest that the existing paradigm of ontogenetic shift in juvenile blue crab habitat utilization requires revision. Although previous evidence supports shifts in blue crab habitat utilization at sizes exceeding 25 mm CW, our results suggest that juvenile blue crabs begin to emigrate from seagrass meadows to salt marsh habitat near 15 mm CW, before progressing to unstructured habitats at larger sizes (i.e. 25–55 mm CW; Lipcius et al., 2005). Emigration to SME at smaller sizes would also explain patterns at larger spatial and temporal scales (Hyman et al., 2022), whereby density of juvenile blue crabs 20–40 mm CW was positively correlated with salt marsh habitat availability, especially in turbid areas. Although low densities of juveniles *>*30 mm CW prevented us from evaluating their habitat use of salt marsh, the present findings and related inferences would benefit from studies that consider additional size classes beyond those included here (e.g. *≤* 15, 16–30, and 31–45 mm CW size classes) to assess whether larger juveniles remain near marsh habitat or emigrate to other unstructured habitats.

### 5.2. Turbidity

In our study, juvenile blue crab abundance was positively associated with turbidity in both size classes. In addition, the association between turbidity and abundance of large juvenile was stronger than that of small juveniles. High turbidity may increase juvenile abundance through both bottom-up and top-down controls. First, turbidity is positively associated with preferred food items of juvenile blue crabs: thin-shelled infaunal bivalves including the soft-shell clam *Mya arenaria* and Baltic clam *Macoma balthica* (Seitz et al., 2003; Seitz, Lipcius and Seebo, 2005). These species constitute a substantial proportion of juvenile blue crab diets and they aggregate near estuarine turbidity maxima within Chesapeake Bay tributaries (Seitz et al., 2003). Turbid upriver unstructured habitats are associated with higher juvenile blue crab growth rates than those in downriver habitats (Seitz, Lipcius and Seebo, 2005). Hence, association between turbidity and juvenile blue crab abundance may be a proxy for high prey abundance and bottom-up control. Second, turbidity may provide protection to juveniles from visual predators through a reduction in detectability (Cyrus and Blaber, 1987; Ajemian, Sohel and Mattila, 2015; Marley et al., 2020), and it may also reduce cannibalism by larger congeners (O’Brien, Slade and Vinyard, 1976). However, many estuarine-dependent predators possess adaptations to forage using chemo-tactile sensors in low-visibility environments characteristic of estuaries, and as a result it is unlikely that high turbidity provides more than a partial refuge from predation (Howson et al., 2022). Whether the association between turbidity and juvenile abundance is due to the former mechanism, the latter, or a combination of both is not addressed here and requires further research.

## 5. Conclusions and future work

Juveniles of marine fish and invertebrates encounter a diverse portfolio of habitats within the estuarine seascape. Habitat characteristics and environmental heterogeneity engender variability in vital rates. The attractiveness of a particular habitat to juveniles is dependent on size or life stage due to changing requirements as juveniles grow (Werner and Gilliam, 1984; Dahlgren and Eggleston, 2000; Lipcius et al., 2005; Johnston and Lipcius, 2012). These shifts in habitat utilization can occur at small sizes (Nagelkerken et al., 2015). The relatively short duration of occupancy, coupled with the tendency of studies to assess juvenile habitat requirements in aggregate (i.e. assessing the needs of immature animals without regard to size-classes), may cause researchers to underestimate the importance of transient habitats essential to juvenile organisms at specific life stages (Nagelkerken et al., 2015). As requirements of juveniles may change most rapidly in their earliest life stages, it is imperative that fine-scale changes in habitat utilization be identified.

Our results both underscore the value of salt marsh habitat for small blue crab juveniles and is consistent with the hypothesis that salt marshes represent a valuable intermediate nursery habitat as larger juveniles move from seagrass meadows to unstructured bottom through ontogeny (Werner and Gilliam, 1984; Dahlgren and Eggleston, 2000; Lipcius et al., 2005). Loss of salt marsh habitat may thus impose a bottleneck in population dynamics as small juveniles emigrate from seagrass beds.

Although juvenile abundance is a key metric when assessing the nursery function of salt marsh habitat, its role in population dynamics requires assessment of secondary production to the adult segment of a population by the use of additional metrics including survival, growth, and juvenile-adult linkage (Beck et al., 2001). For example, high juvenile abundance will not necessarily translate into high secondary production if survival of juveniles to adulthood is low. Further studies using additional metrics, such as growth and survival, concomitantly would help to clarify the role of salt marsh nursery habitats at the population level for blue crabs.

## Acknowledgments

ACH thanks D Eggleston, M Fabrizio, and C Patrick for their ideas as members of ACH’s PhD Committee.

## APPENDIX A: LOGICAL FRAMEWORK

**g_1_**: Abundance is a function of habitat and turbidity. Structurally complex habitats harbor higher densities of juvenile blue crabs relative to unstructured habitats. Hence, relative to sand habitat, juvenile blue crab density is higher seagrass, SME, and SDH Orth and van Montfrans (1987); Heck Jr, Coen and Morgan (2001); Johnson and Eggleston (2010); Etherington and Eggleston (2000); Hyman et al. (2022). Mean-while, high local turbidity increases juvenile abundance through both bottom-up (Seitz et al., 2003; Seitz, Lipcius and Seebo, 2005) and potentially top-down (O’Brien, Slade and Vinyard, 1976; Ajemian, Sohel and Mattila, 2015) mechanisms (see methods in Hyman et al., 2022, for more details).

**g_2_**: Abundance is a function of habitat, turbidity, and an interaction between habitat and turbidity. Here, the effect of turbidity is dependent on a particular habitat. Whereas seagrass meadows are absent from high-turbidity areas due to light requirements, extensive salt marshes and unstructured sand habitats occur in both high- and low-turbidity regions of the tributaries. Turbidity may therefore modify the effectiveness of these habitats as nurseries for juvenile crabs by decreasing predatory foraging efficiency through both low visibility (turbidity) and structural impediments (in SME or SDH; Ajemian, Sohel and Mattila, 2015; Hyman et al., 2022).

**g_3_**: Abundance is a function of habitat, turbidity and spatial position. Recruitment in a given location is dependent on postlarval supply (Beck et al., 2001; Gillanders et al., 2003; Sheaves, Baker and Johnston, 2006). Blue crab postlarvae enter tributaries from the mouth (i.e. downriver), and decline with distance upriver along the tributary axis as they encounter suitable habitat and settle (Stockhausen and Lipcius, 2003). Hence, we expected habitats positioned closer to the mouth of the river would be associated with higher juvenile abundances due to proximity to postlarval supply.

**g_4_**: Abundance is a function of habitat, turbidity, spatial position, and an interaction between habitat and spatial position. Environmental conditions vary substantially along tributary axes (e.g. Posey et al., 2005). Latent variables influencing juvenile abundance may inconsistently affect habitats. As a consequence, the effects of spatial position are habitat-specific (Sheaves et al., 2015; Nagelkerken et al., 2015).

**g_5_**: Abundance is a function of habitat, turbidity, spatial position, and an interaction between habitat and turbidity (Hyman et al., 2022).

**g_6_**: Abundance is a function of habitat, turbidity, spatial position, an interaction between habitat and spatial position, and an interaction between habitat and turbidity. Additional environmental variables other than turbidity may augment habitat suitability at different spatial positions, such as salinity (e.g. Posey et al., 2005) and/or food availability (e.g. Seitz, Lipcius and Seebo, 2005).

## APPENDIX B: PRIOR DISTRIBUTIONS FOR GEAR EFFICIENCY

As different sampling methods were employed for the four habitat types, gear efficiency estimates were required to scale abundance estimates for each sample. Efficiency of the suction sampling methodology is estimated at 0.88 Orth and van Montfrans (1987). Meanwhile, pilot efficiency tests of the modified flume net design using marked blue crabs in fall of 2020 indicated an estimated efficiency of 0.92 *±* 0.02. Finally, juvenile blue crab depletion experiments for benthic scrape gear suggested efficiency between 0.21 and 0.45 (Ralph and Lipcius, 2014). However, benthic scrapes used here differed slightly in that they did not include iron teeth, which may decrease efficiency.

We constructed normally distributed prior distributions for each gear type based on estimates from literature and observed data and subsequently applied a log-transformation to relate prior estimates of efficiency to the model scale. For flume traps, the prior distribution was ln *N* (0.92, 0.02) which has a mean of-0.083 and standard deviation of 0.02. Similarly, for scrape estimates, we assumed the mean efficiency was 0.33 (average of 0.45 and 0.21) and a standard deviation of 0.12 to yield a prior of ln *N* (0.33, 0.12) which has a mean of -1.2 and standard deviation of 0.18. Finally, for suction sampling, average efficiency is 0.88 (Orth and van Montfrans, 1987), although uncertainty estimates were not supplied in literature. Here, we assumed an efficiency of 0.02 similar to uncertainty in flume traps and applied a prior of ln *N* (0.88, 0.02) which has a mean of -0.13 and standard deviation of 0.02.

## APPENDIX C: SUPPLEMENTAL TABLES

**TABLE S1.** Data summaries of crab carapace widths (CW) and physicochemical variables.

|  | Temperature (°C) | Salinity | Secchi (cm) | DO (mg/L) | CW (mm) |
| --- | --- | --- | --- | --- | --- |
| Min | 17.30 | 2.66 | 8.20 | 3.90 | 1.00 |
| 10% | 19.90 | 6.06 | 30.00 | 6.28 | 6.40 |
| 50% | 24.10 | 17.56 | 78.00 | 7.63 | 12.90 |
| 90% | 28.90 | 19.46 | 128.41 | 8.77 | 25.28 |
| Max | 30.90 | 20.32 | 183.00 | 11.37 | 135.10 |

**TABLE S2.** Table displaying the number of samples for each habitat by trip. Five of the total 144 samples were expunged due to missing predictor values (i.e. Secchi disk depth) in seagrass (two samples) and SDH (three samples)

| Habitat | Trip 1 | Trip 2 | Trip 3 | Trip 4 | Total |
| --- | --- | --- | --- | --- | --- |
| SME | 0 | 6 | 6 | 6 | 18 |
| SDH | 15 | 18 | 18 | 0 | 51 |
| Seagrass | 4 | 6 | 6 | 0 | 16 |
| Sand | 18 | 18 | 18 | 0 | 54 |

## APPENDIX D: SUPPLEMENTARY FIGURES

**Fig S1.**
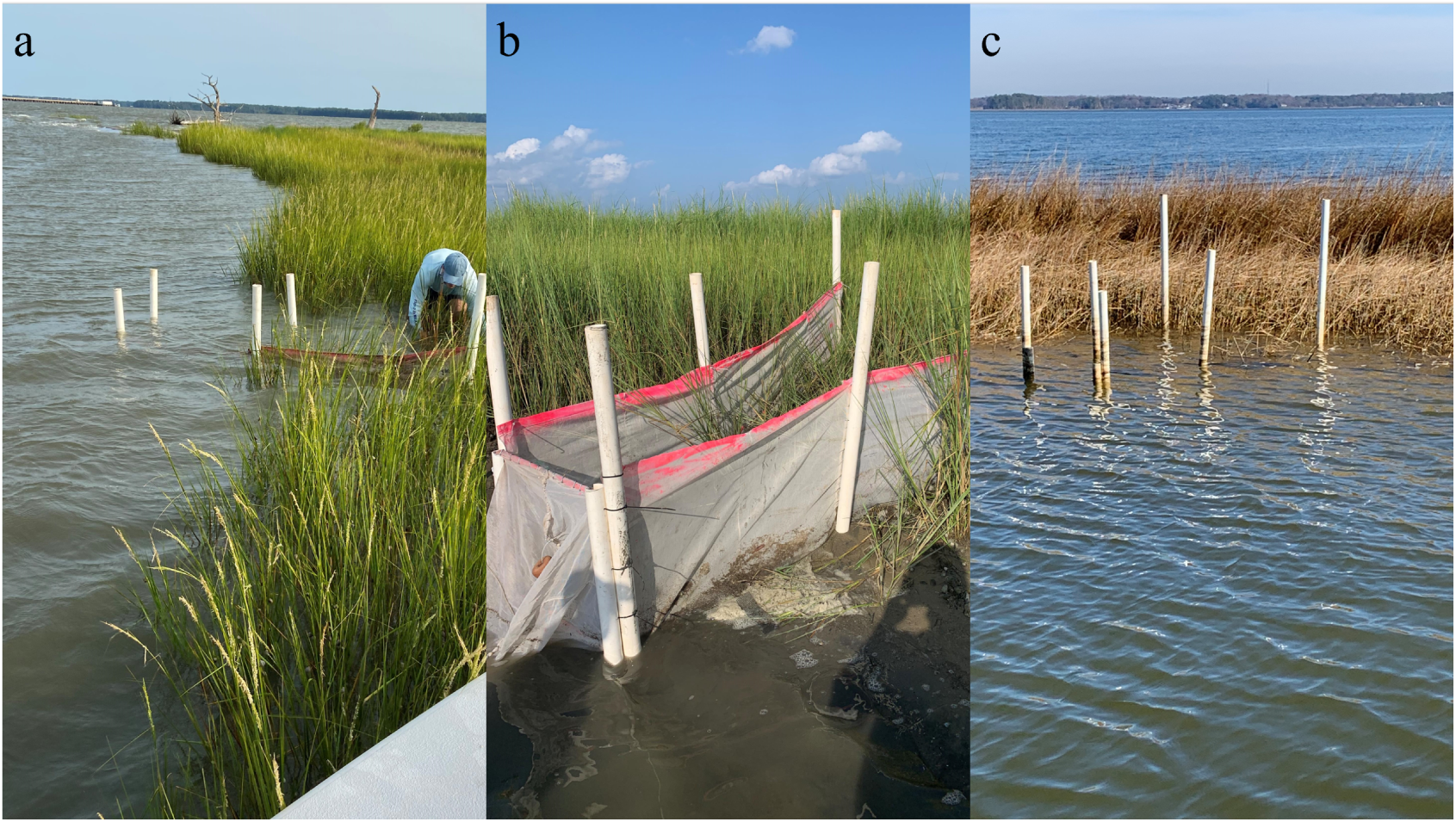
Images of flume net in multiple stages of deployment: **a)** depicts a flume net set up at slack flood tide; **b)** denotes flume net collected at slack ebb tide; and **c)** denotes flume in non-deployment stage with net walls down and end removed when net is not in use. When not in use, enclosures will remain on site with the net walls folded and staked into the ground and the end removed, facilitating movement of animals throughout marsh habitat. Prior to use, walls of the flume nets will be rapidly erected to contain all animals occupying the habitat at the time of sampling.

**Fig S2.**
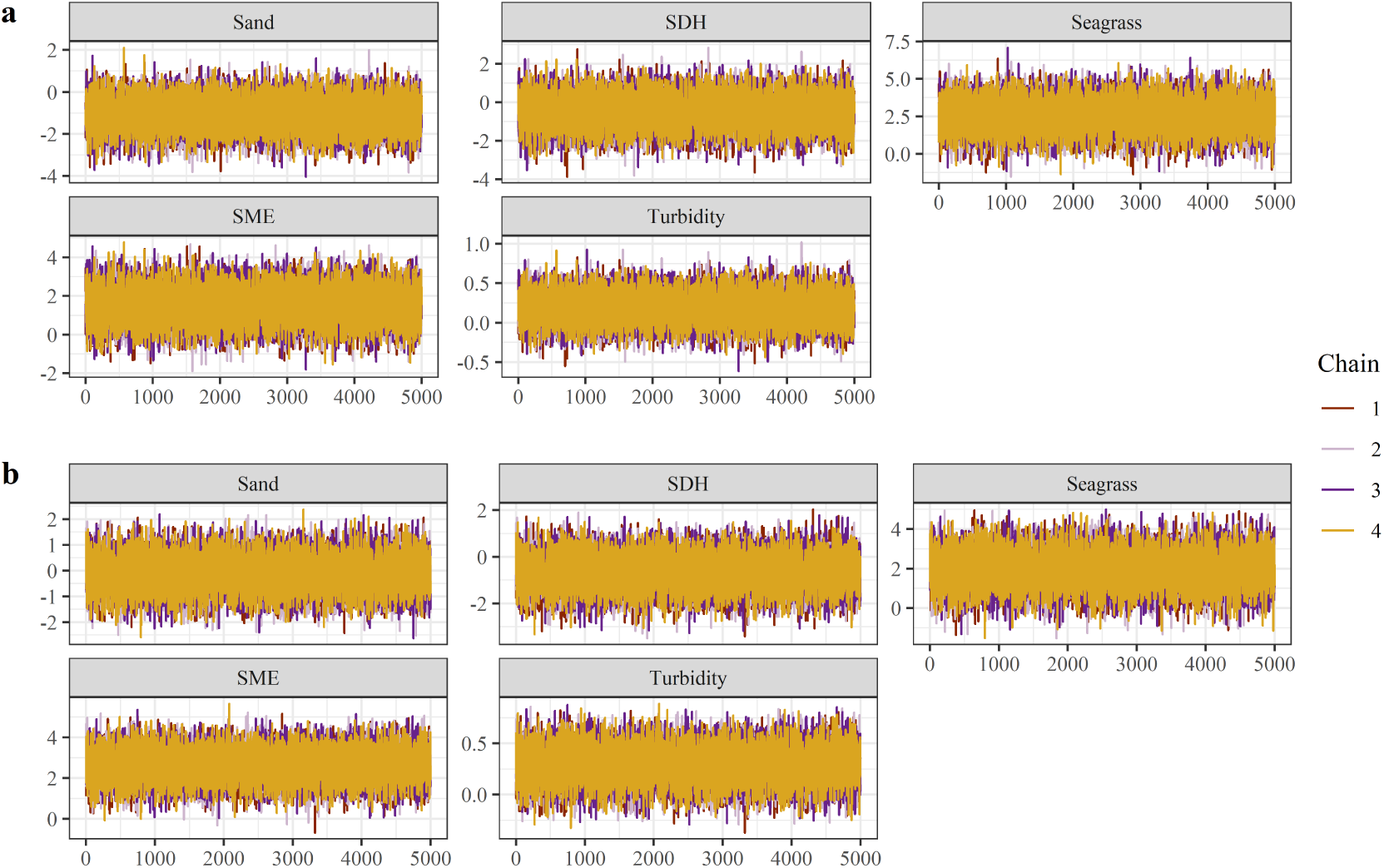
A set of trace plots for model g_1_ parameters illustrating posterior values of each regression coefficient per Markov chain throughout the post-warmup/adaptive phase. Visual inspection of trace plots is used to evaluate convergence and mixing of the chains.

**Fig S3.**
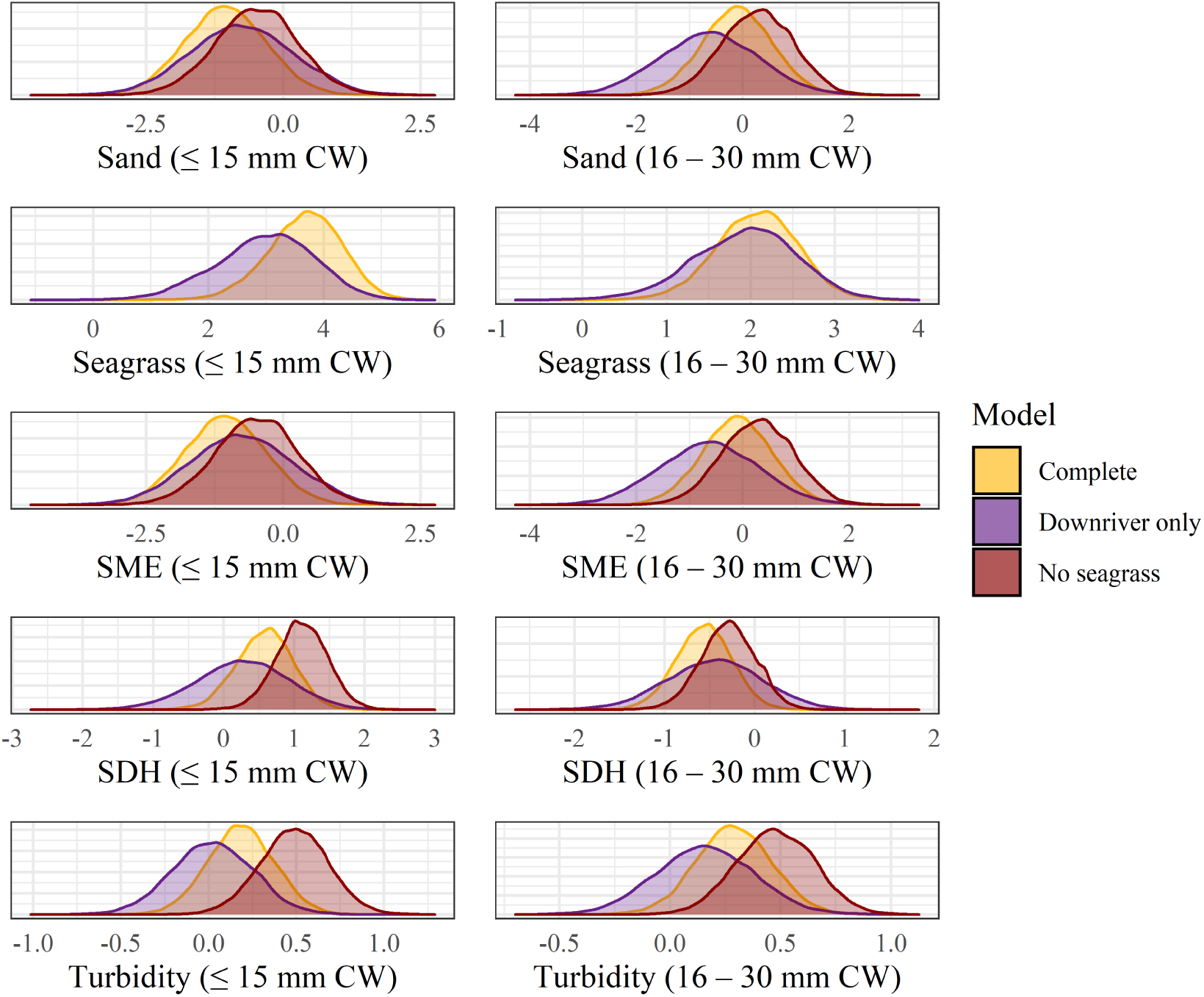
Posterior distributions of regression coefficients from the selected model g_1_ sing the complete data set (Complete, yellow), and subsets of the data using only the downriver stratum (Downriver only, purple) and without seagrass (No seagrass, maroon). Posterior distributions were largely consistent across models, indicating inferences on the complete data set were robust.

**Fig S4.**
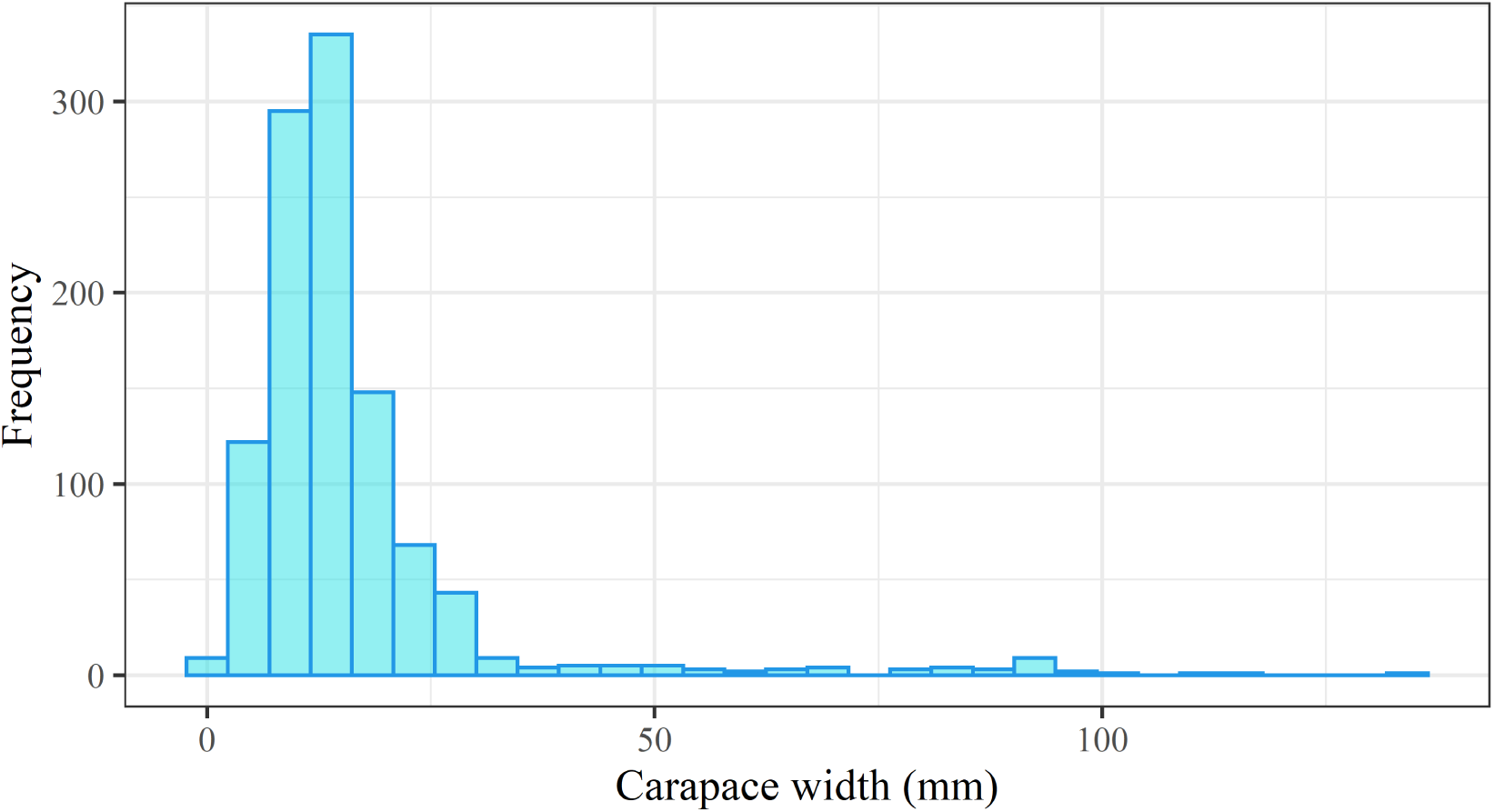
Data histogram of all crab carapace widths (mm) caught in Fall 2020 recruitment period.

**Fig S5.**
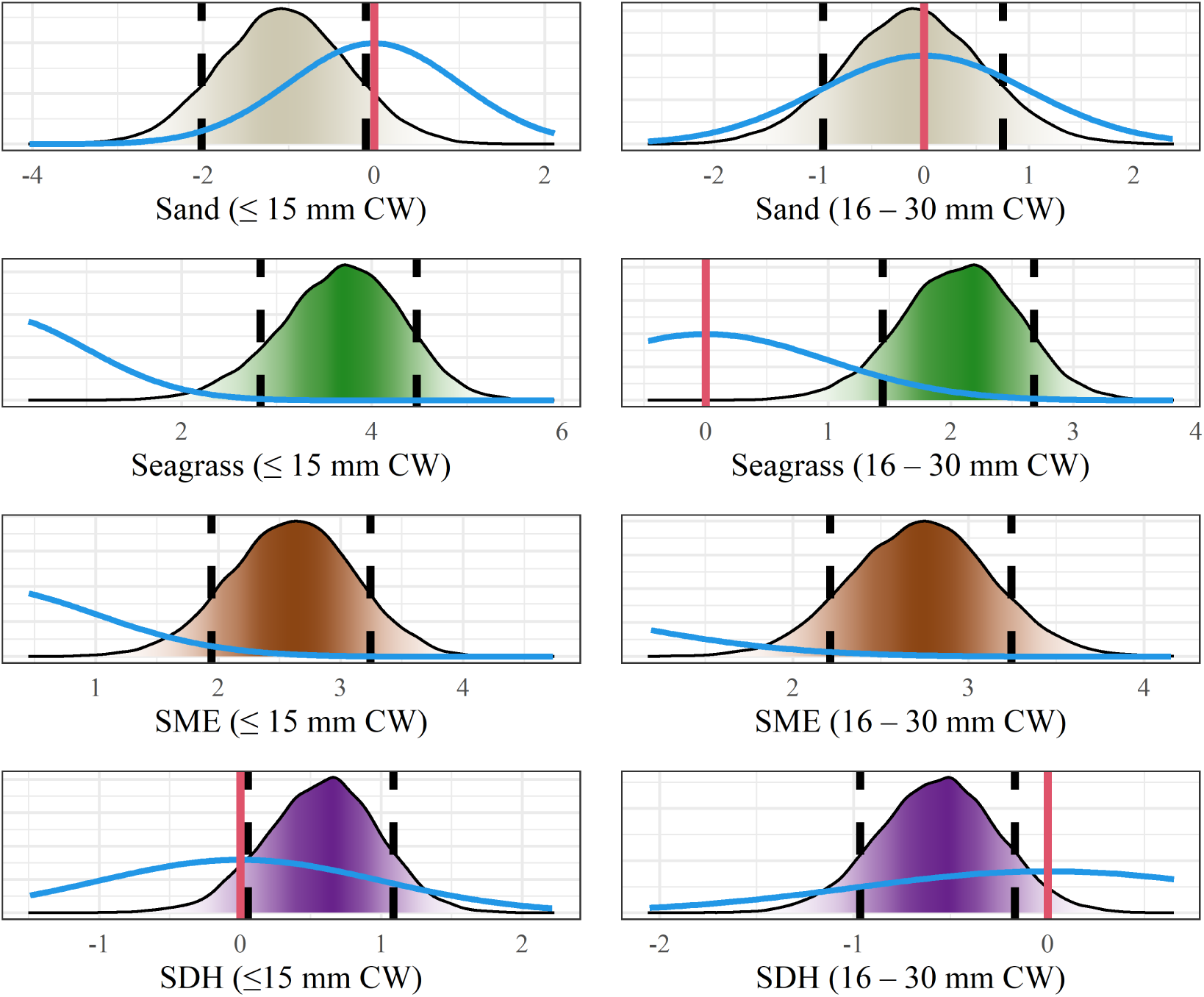
*Conditional posterior distributions of* ln *mean habitat-specific abundances (conditioned on holding* ln *turbidity at 0) from model g*_1_ *for both small (≤ 15 mm CW; left column) and large (16–30 mm CW; right column) size classes. Dashed black lines denote 80% Bayesian confidence intervals, while red lines (where present) denote 0. Blue lines depict prior distributions*.

**Fig S6.**
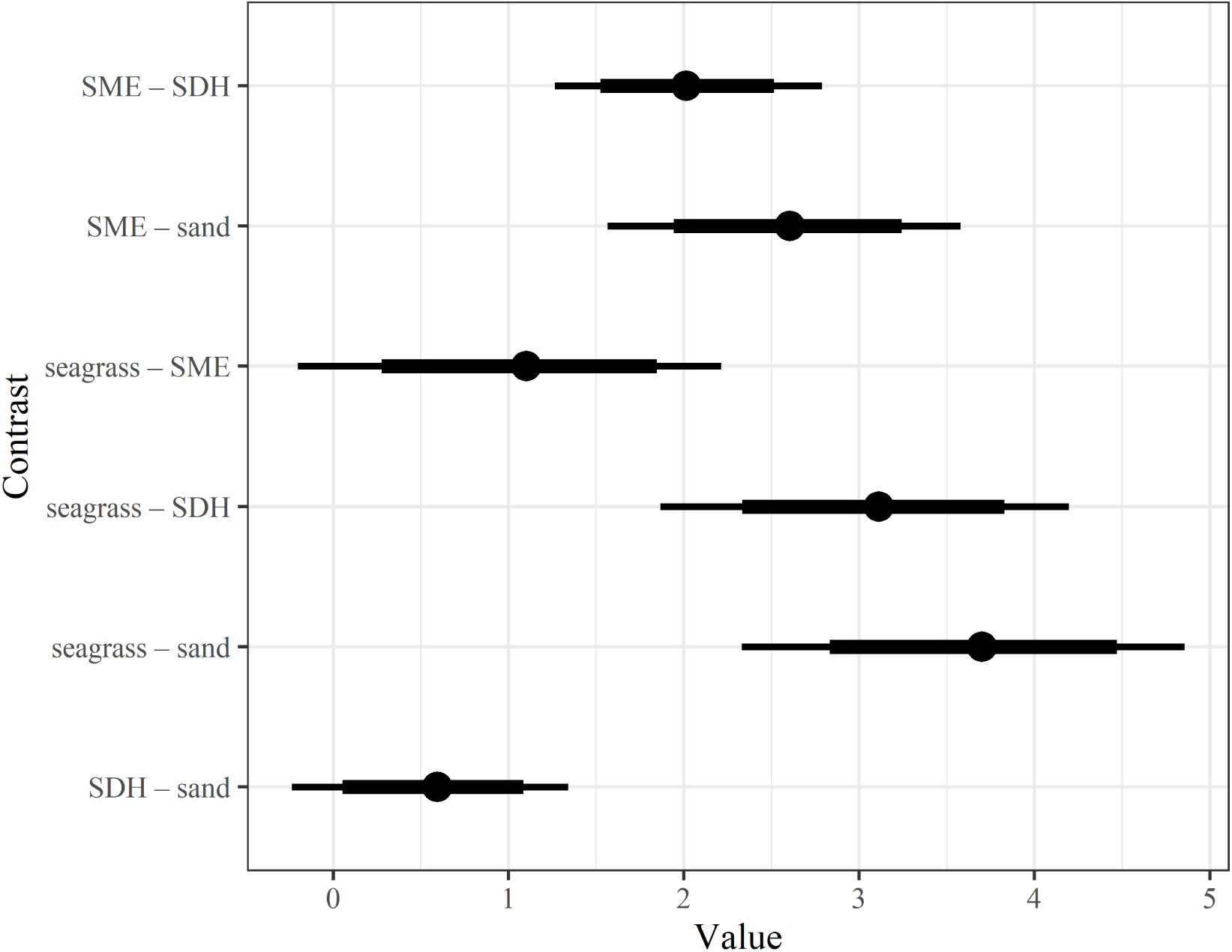
*Linear contrast statements (see Section 3.3.3) depicting differences in* ln *expected juvenile blue crab abundance for the small size class from Model g*_1_*. Dots denote posterior median difference in* ln *expected values, while thick bars represent 80% Bayesian CI’s and thin bars denote 95% Bayesian CI’s*.

**Fig S7.**
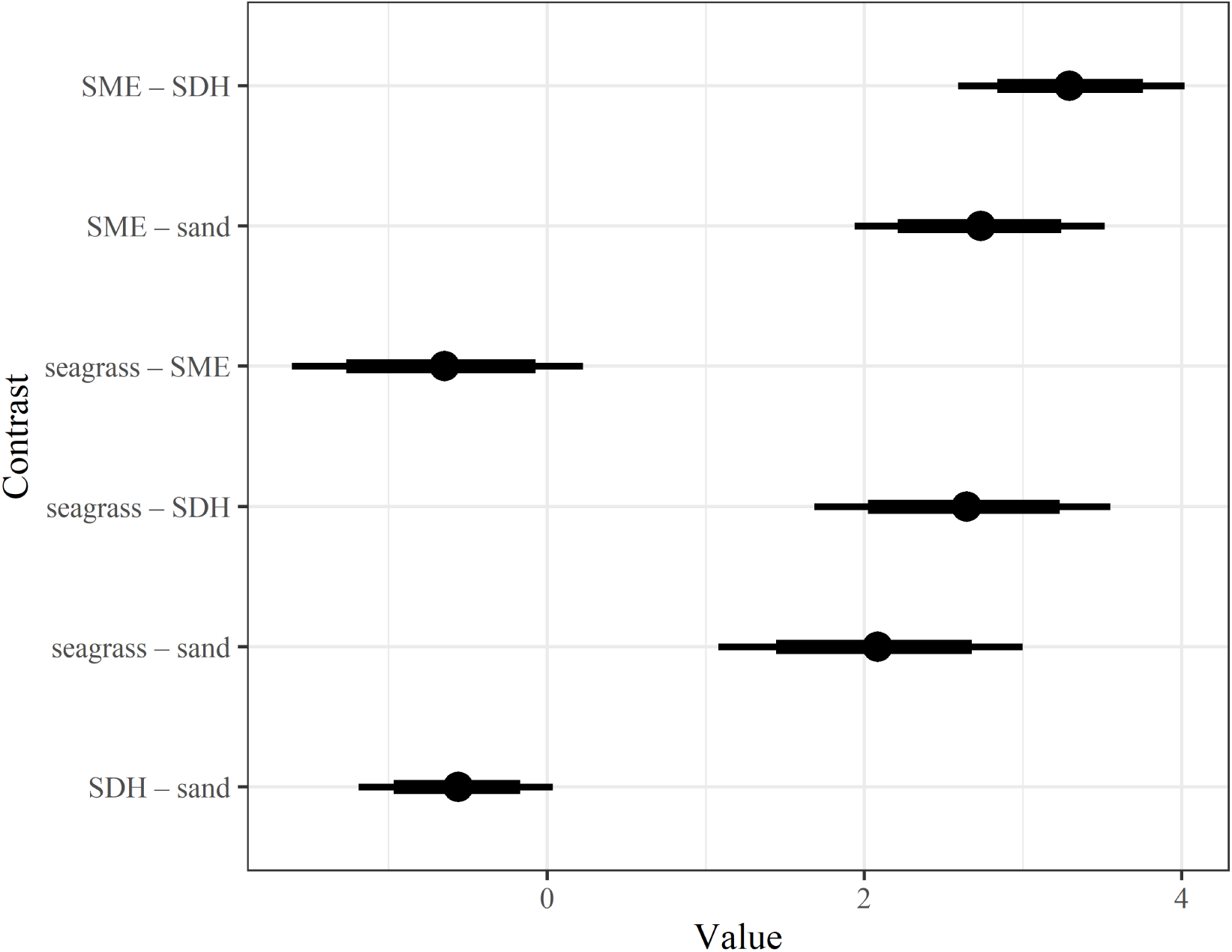
*Linear contrast statements (see Section 3.3.3) depicting differences in* ln *expected juvenile blue crab abundance for the large size class from Model g*_1_*. Dots denote posterior median differences in* ln *expected values, while thick bars represent 80% Bayesian CI’s and thin bars denote 95% Bayesian CI’s*.

**Fig S8.**
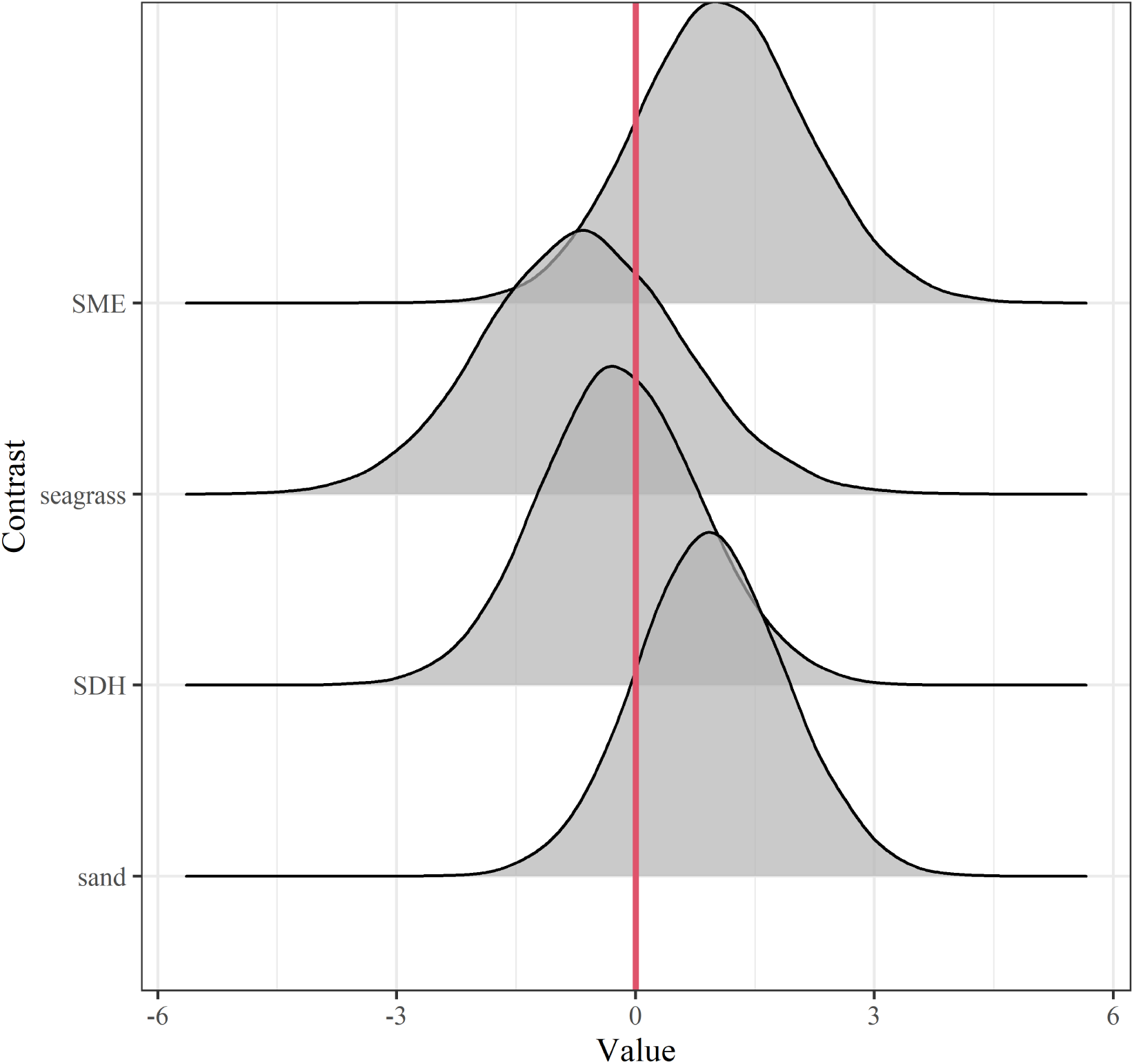
*Posterior distributions of within-habitat linear contrasts (see Section 3.3.3) depicting differences in* ln *expected juvenile blue crab abundance for small and large size class from Model g*_1_*. Positive values indicate increases in expected abundance as animals move from ≤ 15 to 16–30 mm, while negative values indicate decreases in expected abundance*.

## REFERENCES

Aburto-Oropeza, O., Dominguez-Guerrero, I., Cota-Nieto, J. and Plomozo-Lugo, T. (2009). Recruitment and ontogenetic habitat shifts of the yellow snapper (*Lutjanus argentiventris*) in the Gulf of California. Marine Biology 156 2461–2472.

Adams, A. J., Dahlgren, C. P., Kellison, G. T., Kendall, M. S., Layman, C. A., Ley, J. A., Nagelkerken, I. and Serafy, J. E. (2006). Nursery function of tropical back-reef systems. Marine Ecology Progress Series 318 287–301.

Ajemian, M., Sohel, S. and Mattila, J. (2015). Effects of turbidity and habitat complexity on antipredator behavior of three-spined sticklebacks (*Gasterosteus aculeatus*). Environmental Biology of Fishes 98 45–55.

Amorim, E., Ramos, S., Elliott, M. and Bordalo, A. A. (2018). Dynamic habitat use of an estuarine nursery seascape: Ontogenetic shifts in habitat suitability of the European flounder (*Platichthys flesus*). Journal of Experimental Marine Biology and Ecology 506 49–60.

Beck, M. W., Heck, K. L., Able, K. W., Childers, D. L., Eggleston, D. B., Gillanders, B. M., Halpern, B., Hays, C. G., Hoshino, K., Minello, T. J., Orth, R. J., Sheridan, P. F. and Weinstein, M. P. (2001). The identification, conservation, and management of estuarine and marine nurseries for fish and invertebrates: a better understanding of the habitats that serve as nurseries for marine species and the factors that create site-specific variability in nursery quality will improve conservation and management of these areas. Bioscience 51 633–641.

Berkström, C., Lindborg, R., Thyresson, M. and GullströM, M. (2013). Assessing connectivity in a tropical embayment: fish migrations and seascape ecology. Biological Conservation 166 43–53.

Bishop, T. D., Miller, H. L., Walker, R. L., Hurley, D. H., Menken, T. and Tilburg, C. E. (2010). Blue crab (Callinectes sapidus Rathbun, 1896) settlement at three Georgia (USA) estuarine sites. Estuaries and Coasts 33 688–698.

Blackmon, D. C. and Eggleston, D. B. (2001). Factors influencing planktonic, post-settlement dispersal of early juvenile blue crabs (*Callinectes sapidus* Rathbun). Journal of Experimental Marine Biology and Ecology 257 183–203.

Cicchetti, G. (1998). Habitat use, secondary production, and trophic export by salt marsh nekton in shallow waters, PhD thesis, The College of William and Mary.

Cicchetti, G. and Diaz, R. (2000). Types of salt marsh edge and export of trophic energy from marshes to deeper habitats. In Concepts and controversies in tidal marsh ecology (M. P. Weinstein and D. A. Kreeger, eds.) 535–564. Kluwer.

Ciotti, B., Brown, E., Colloca, F., Le Pape, O., Hyman, A. C., Lipcius, R. N., Seitz, R., Maathuis, M., Poiesz, S., Rose, K., Ventura, D. and Van De Wolfshaar, K. (in prep). Measuring the quality of nursery habitats for juvenile fish and invertebrates: perspectives from 50 years of research.

Clark, K. L., Ruiz, G. M. and Hines, A. H. (2003). Diel variation in predator abundance, predation risk and prey distribution in shallow-water estuarine habitats. Journal of Experimental Marine Biology and Ecology 287 37–55.

Crowder, L. B. and Cooper, W. E. (1982). Habitat structural complexity and the interaction between bluegills and their prey. Ecology 63 1802–1813.

Crowder, L. B., Squires, D. D. and Rice, J. A. (1997). Nonadditive effects of terrestrial and aquatic predators on juvenile estuarine fish. Ecology 78 1796–1804.

Cyrus, D. and Blaber, S. (1987). The influence of turbidity on juvenile marine fish in the estuaries of Natal, South Africa. Continental Shelf Research 7 1411–1416.

Dahlgren, C. P. and Eggleston, D. B. (2000). Ecological processes underlying ontogenetic habitat shifts in a coral reef fish. Ecology 81 2227–2240.

Dahlgren, C. P., Kellison, G. T., Adams, A. J., Gillanders, B. M., Kendall, M. S., Layman, C. A., Ley, J. A., Nagelkerken, I. and Serafy, J. E. (2006). Marine nurseries and effective juvenile habitats: concepts and applications. Marine Ecology Progress Series 312 291–295.

Dorenbosch, M., Grol, M. G., De Groene, A., Van Der Velde, G. and Nagelkerken, I. (2009). Piscivore assemblages and predation pressure affect relative safety of some back-reef habitats for juvenile fish in a Caribbean bay. Marine Ecology Progress Series 379 181–196.

Epifanio, C. (2007). Biology of larvae. *The Blue Crab, Callinectes sapidus. Maryland Sea Grant, College Park*, MD 513–533.

Epifanio, C. E. (2019). Early life history of the blue crab Callinectes sapidus: a review. Journal of Shellfish Research 38 1–22.

Etherington, L. L. and Eggleston, D. B. (2000). Large-scale blue crab recruitment: linking postlarval transport, post-settlement planktonic dispersal, and multiple nursery habitats. Marine Ecology Progress Series 204 179–198.

Etherington, L. L., Eggleston, D. B. and Stockhausen, W. T. (2003). Partitioning loss rates of early juvenile blue crabs from seagrass habitats into mortality and emigration. Bulletin of Marine Science 72 371–391.

Fitz, H. C. and Wiegert, R. G. (1991). Utilization of the intertidal zone of a salt marsh by the blue crab *Callinectes sapidus*: density, return frequency, and feeding habits. Marine Ecology Progress Series 249–260.

Forrester, G. E. and Swearer, S. E. (2002). Trace elements in otoliths indicate the use of open-coast versus bay nursery habitats by juvenile California halibut. Marine Ecology Progress Series 241 201–213.

Fraser, D. F. and Emmons, E. E. (1984). Behavioral response of blacknose dace (*Rhinichthys atratulus*) to varying densities of predatory creek chub (*Semotilus atromaculatus*). Canadian Journal of Fisheries and Aquatic Sciences 41 364–370.

Gelman, A., Lee, D. and Guo, J. (2015). Stan: a probabilistic programming language for Bayesian inference and optimization. Journal of Educational and Behavioral Statistics 40 530–543.

Gillanders, B. M., Able, K. W., Brown, J. A., Eggleston, D. B. and Sheridan, P. F. (2003). Evidence of connectivity between juvenile and adult habitats for mobile marine fauna: an important component of nurseries. Marine Ecology Progress Series 247 281–295.

Heck Jr, K., Coen, L. and Morgan, S. (2001). Pre- and post-settlement factors as determinants of juvenile blue crab *Callinectes sapidus* abundance: results from the north-central Gulf of Mexico. Marine Ecology Progress Series 222 163–176.

Heck Jr, K., Hays, G. and Orth, R. J. (2003). Critical evaluation of the nursery role hypothesis for seagrass meadows. Marine Ecology Progress Series 253 123–136.

Hovel, K. A. and Fonseca, M. S. (2005). Influence of seagrass landscape structure on the juvenile blue crab habitat-survival function. Marine Ecology Progress Series 300 179–191.

Hovel, K. A. and Lipcius, R. N. (2002). Effects of seagrass habitat fragmentation on juvenile blue crab survival and abundance. Journal of Experimental Marine Biology and Ecology 271 75–98.

Howson, U. A. (2000). Nursery habitat quality for juvenile paralichthyid flounders: Experiemental analyses of the effects of physicochemical parameters, PhD thesis, University of Delaware.

Howson, U. A., Targett, T. E., Grecay, P. A. and Gaffney, P. M. (2022). Foraging by estuarine juveniles of two *paralichthyid* flounders: experimental analyses of the effects of light level, turbidity, and prey type. Marine Ecology Progress Series 695 139–156.

Hyman, A. C., Chiu, G. S., Fabrizio, M. C. and Lipcius, R. N. (2022). Spatiotemporal Modeling of Nursery Habitat Using Bayesian Inference: Environmental Drivers of Juvenile Blue Crab Abundance. Frontiers in Marine Science 9 834990.

Isdell, R. E., Bilkovic, D. M., Guthrie, A. G., Mitchell, M. M., Chambers, R. M., Leu, M. and Hershner, C. (2021). Living shorelines achieve functional equivalence to natural fringe marshes across multiple ecological metrics. PeerJ 9 e11815.

Jivoff, P. R. and Able, K. W. (2003). Evaluating salt marsh restoration in Delaware Bay: the response of blue crabs, *Callinectes sapidus*, at former salt hay farms. Estuaries 26 709–719.

Johnson, E. G. and Eggleston, D. B. (2010). Population density, survival and movement of blue crabs in estuarine salt marsh nurseries. Marine Ecology Progress Series 407 135–147.

Johnston, C. A. and Caretti, O. N. (2017). Mangrove expansion into temperate marshes alters habitat quality for recruiting *Callinectes spp*. Marine Ecology Progress Series 573 1–14.

Johnston, C. A. and Lipcius, R. N. (2012). Exotic macroalga *Gracilaria vermiculophylla* provides superior nursery habitat for native blue crab in Chesapeake Bay. Marine Ecology Progress Series 467 137–146.

Jones, D. L., Walter, J. F., Brooks, E. N. and Serafy, J. E. (2010). Connectivity through ontogeny: fish population linkages among mangrove and coral reef habitats. Marine Ecology Progress Series 401 245–258.

Kruschke, J. K. (2021). Bayesian analysis reporting guidelines. Nature Human Behaviour 5 1282–1291.

Lefcheck, J. S., Hughes, B. B., Johnson, A. J., Pfirrmann, B. W., Rasher, D. B., Smyth, A. R., Williams, B. L., Beck, M. W. and Orth, R. J. (2019). Are coastal habitats important nurseries? A meta-analysis. Conservation Letters 12 e12645.

Lipcius, R. N., Seitz, R. D., Seebo, M. S. and Colón-CarrióN, D. (2005). Density, abundance and survival of the blue crab in seagrass and unstructured salt marsh nurseries of Chesapeake Bay. Journal of Experimental Marine Biology and Ecology 319 69–80.

Lipcius, R. N., Eggleston, D. B., Heck Jr, K. L., Seitz, R. D. and Van Montrans, J. (2007). Post-settlement abundance, survival, and growth of postlarvae and young juvenile blue crabs in nursery habitats. *The Blue Crab Callinectes sapidus. Maryland Sea Grant College, College Park*, Maryland 535–564.

Lipcius, R. N., Eggleston, D. B., Fodrie, F. J., Van Der Meer, J., Rose, K. A., Vasconcelos, R. P. and Van De Wolfshaar, K. E. (2019). Modeling quantitative value of habitats for marine and estuarine populations. Frontiers in Marine Science 6 280.

Litvin, S. Y., Weinstein, M. P., Sheaves, M. and Nagelkerken, I. (2018). What makes nearshore habitats nurseries for nekton? An emerging view of the nursery role hypothesis. Estuaries and Coasts 41 1539–1550.

Lugendo, B. R., Pronker, A., Cornelissen, I., De Groene, A., Nagelkerken, I., Dorenbosch, M., Van Der Velde, G. And Mgaya, Y. D. (2005). Habitat utilisation by juveniles of commercially important fish species in a marine embayment in Zanzibar, Tanzania. Aquatic Living Resources 18 149–158.

Marley, G. S., Deacon, A. E., Phillip, D. A. and Lawrence, A. J. (2020). Mangrove or mudflat: prioritizing fish habitat for conservation in a turbid tropical estuary. Estuarine, Coastal and Shelf Science 240 106788.

Mcelreath, R. (2018). Statistical rethinking: a Bayesian course with examples in R and Stan. Chapman and Hall/CRC.

Mcivor, C. C. and Odum, W. E. (1986). The flume net: a quantitative method for sampling fishes and macrocrustaceans on tidal marsh surfaces. Estuaries 9 219–224.

Miller, C. R., Hyman, A. C., Shi, D. and Lipcius, R. N. (2023). Test of treatment-specific bias in simulated marsh with juvenile blue crab *Callinectes sapidus*. In prep.

Minello, T. J., Able, K. W., Weinstein, M. P. and Hays, C. G. (2003). Salt marshes as nurseries for nekton: testing hypotheses on density, growth and survival through meta-analysis. Marine Ecology Progress Series 246 39–59.

Mizerek, T., Regan, H. M. and Hovel, K. A. (2011). Seagrass habitat loss and fragmentation influence management strategies for a blue crab *Callinectes sapidus* fishery. Marine Ecology Progress Series 427 247–257.

Nagelkerken, I., Sheaves, M., Baker, R. and Connolly, R. M. (2015). The seascape nursery: a novel spatial approach to identify and manage nurseries for coastal marine fauna. Fish and Fisheries 16 362–371.

Nakamura, Y., Hirota, K., Shibuno, T. and Watanabe, Y. (2012). Variability in nursery function of tropical seagrass beds during fish ontogeny: timing of ontogenetic habitat shift. Marine Biology 159 1305–1315.

O’Brien, W. J., Slade, N. A. and Vinyard, G. L. (1976). Apparent size as the determinant of prey selection by bluegill sunfish (*Lepomis macrochirus*). Ecology 57 1304–1310.

Orth, R. J. and Van Montfrans, J. (1987). Utilization of a seagrass meadow and tidal marsh creek by blue crabs *Callinectes sapidus*. Seasonal and annual variations in abundance with emphasis on post-settlement juveniles. Marine Ecology Progress Series 41 283.

Orth, R. J. and Van Montfrans, J. (2002). Habitat quality and prey size as determinants of survival in post-larval and early juvenile instars of the blue crab *Callinectes sapidus*. Marine Ecology Progress Series 231 205–213.

Perkins-Visser, E., Wolcott, T. G. and Wolcott, D. L. (1996). Nursery role of seagrass beds: enhanced growth of juvenile blue crabs (*Callinectes sapidus Rathbun*). Journal of Experimental Marine Biology and Ecology 198 155–173.

Peterson, C. H. and Lipcius, R. N. (2003). Conceptual progress towards predicting quantitative ecosystem benefits of ecological restorations. Marine Ecology Progress Series 264 297–307.

Pile, A. J., Lipcius, R. N., Van Montfrans, J. and Orth, R. J. (1996). Density-dependent settler-recruit-juvenile relationships in blue crabs. Ecological Monographs 66 277–300.

Posey, M. H., Alphin, T. D., Harwell, H. and Allen, B. (2005). Importance of low salinity areas for juvenile blue crabs, *Callinectes sapidus Rathbun*, in river-dominated estuaries of southeastern United States. Journal of Experimental Marine Biology and Ecology 319 81–100.

Ralph, G. (2014). Quantification of Nursery Habitats for Blue Crabs in Chesapeake Bay, PhD thesis, Virginia Institute of Marine Science, William & Mary.

Ralph, G. M. and Lipcius, R. N. (2014). Critical habitats and stock assessment: age-specific bias in the Chesapeake Bay blue crab population survey. Transactions of the American Fisheries Society 143 889–898.

Ralph, G. M., Seitz, R. D., Orth, R. J., Knick, K. E. and Lipcius, R. N. (2013). Broad-scale association between seagrass cover and juvenile blue crab density in Chesapeake Bay. Marine Ecology Progress Series 488 51–63.

Reyns, N. B. and Eggleston, D. B. (2004). Environmentally-controlled, density-dependent secondary dispersal in a local estuarine crab population. Oecologia 140 280–288.

Rozas, L. P. and Minello, T. J. (1998). Nekton use of salt marsh, seagrass, and nonvegetated habitats in a south Texas (USA) estuary. Bulletin of Marine Science 63 481–501.

Seitz, R. D., Lipcius, R. N. and Seebo, M. S. (2005). Food availability and growth of the blue crab in seagrass and unvegetated nurseries of Chesapeake Bay. Journal of Experimental Marine Biology and Ecology 319 57–68.

Seitz, R. D., Lipcius, R. N., Stockhausen, W. T., Delano, K. A., Seebo, M. S. and Gerdes, P. D. (2003). Potential bottom-up control of blue crab distribution at various spatial scales. Bulletin of Marine Science 72 471–490.

Seitz, R. D., Wennhage, H., BergströM, U., Lipcius, R. N. and Ysebaert, T. (2014). Ecological value of coastal habitats for commercially and ecologically important species. ICES Journal of Marine Science 71 648–665.

Shakeri, L. M., Darnell, K. M., Carruthers, T. J. and Darnell, M. Z. (2020). Blue crab abundance and survival in a fragmenting coastal marsh system. Estuaries and Coasts 43 1545–1555.

Sheaves, M., Baker, R. and Johnston, R. (2006). Marine nurseries and effective juvenile habitats: an alternative view. Marine Ecology Progress Series 318 303–306.

Sheaves, M., Baker, R., Nagelkerken, I. and Connolly, R. M. (2015). True value of estuarine and coastal nurseries for fish: incorporating complexity and dynamics. Estuaries and Coasts 38 401–414.

Sivula, T., Magnusson, M., Matamoros, A. A. and Vehtari, A. (2020). Uncertainty in Bayesian leave-one-out cross-validation based model comparison. arXiv preprint arXiv:2008.10296.

Smock, L. A., Wright, A. B. and Benke, A. C. (2005). Atlantic coast rivers of the southeastern United States. Rivers of North America 72 122.

Stockhausen, W. T. and Lipcius, R. N. (2003). Simulated effects of seagrass loss and restoration on settlement and recruitment of blue crab postlarvae and juveniles in the York River, Chesapeake Bay. Bulletin of Marine Science 72 409–422.

R Core Team (2022). R: A Language and Environment for Statistical Computing R Foundation for Statistical Computing, Vienna, Austria. 30

Thomas, J., Zimmerman, R. and Minello, T. (1990a). Abundance patterns of juvenile blue crabs (*Callinectes sapidus*) in nursery habitats of two Texas bays. Bulletin of Marine Science 46 115–125.

Thomas, J., Zimmerman, R. and Minello, T. (1990b). Abundance patterns of juvenile blue crabs (*Callinectes sapidus*) in nursery habitats of two Texas bays. Bulletin of Marine Science 46 115–125.

Underwood, A., Chapman, M. and Connell, S. (2000). Observations in ecology: you can’t make progress on processes without understanding the patterns. Journal of Experimental Marine Biology and Ecology 250 97–115.

Urban, M. C. (2007). The growth–predation risk trade-off under a growing gape-limited predation threat. Ecology 88 2587– 2597.

Van Montfrans, J., Ryer, C. H. and Orth, R. J. (2003). Substrate selection by blue crab *Callinectes sapidus* megalopae and first juvenile instars. Marine Ecology Progress Series 260 209–217.

Vasconcelos, R. P., Eggleston, D. B., Le Pape, O. and Tulp, I. (2014). Patterns and processes of habitat-specific demographic variability in exploited marine species. ICES Journal of Marine Science 71 638–647.

Vehtari, A., Gelman, A. and Gabry, J. (2017). Practical Bayesian model evaluation using leave-one-out cross-validation and WAIC. Statistics and Computing 27 1413–1432.

Vehtari, A., Gabry, J., Magnusson, M., Yao, Y., BÜRKNER, P.-C., Paananen, T. and Gelman, A. (2022). loo: Efficient leave-one-out cross-validation and WAIC for Bayesian models. R package version 2.5.1.

Voigt, E. P. and Eggleston, D. B. (2022). Spatial Variation in Nursery Habitat Use by Juvenile Blue Crabs in a Shallow, Wind-Driven Estuary. Estuaries and Coasts 1–15.

Watanabe, S. (2013). A widely applicable Bayesian information criterion. The Journal of Machine Learning Research 14 867–897.

Werner, E. E. and Gilliam, J. F. (1984). The ontogenetic niche and species interactions in size-structured populations. Annual review of ecology and systematics 15 393–425.

Wood, M. A. and Lipcius, R. N. (2022). Non-native red alga *Gracilaria vermiculophylla* compensates for seagrass loss as blue crab nursery habitat in the emerging Chesapeake Bay ecosystem. PloS One 17 e0267880.

Zimmerman, R. J., Minello, T. J. and Rozas, L. P. (2000). Salt marsh linkages to productivity of penaeid shrimps and blue crabs in the northern Gulf of Mexico. 293–314.

